# Siphonous green macroalgae with contrasting capacities for the energy-dependent quenching, qE, rely on different photoprotective mechanisms

**DOI:** 10.1101/2025.08.23.671893

**Authors:** Heta Mattila, Vesa Havurinne, Paulo Cartaxana, Sónia Cruz

## Abstract

To combat light-induced damage, photosynthetic organisms have evolved several photoprotective mechanisms, including non-photochemical quenching (NPQ) of excess light energy. Bryopsidales green macroalgae lack the energy-dependent quenching (qE) and can therefore induce NPQ only slowly. We illuminated a Bryopsidales macroalga (*Bryopsis* sp.) and a morphologically similar Dasycladales alga (*Acetabularia acetabulum*), capable of qE, with high light. No differences in the rate of Photosystem II (PSII) damage, probed by the chlorophyll *a* fluorescence parameter F_V_/F_M_ in the absence and presence of lincomycin, were observed between constant and fluctuating light in *Bryopsis* sp., nor between the two algae. In *Bryopsis* sp., however, photoinhibition lead to decreased rates of electron transfer (estimated by fluorescence and oxygen evolution), while a stimulation was observed in *A. acetabulum*. Compared to *A. acetabulum*, *Bryopsis* sp. showed slow PSII repair, but as the rate of repair increased with increasing rates of the damage, the slow repair may not indicate increased oxidative stress but be a regulatory response. We observed, in *A. acetabulum*, a post-illumination dip in PSII activity and a concurring bump in NPQ, which were removed by an inhibitor of the NDA2-dependent cyclic electron transfer route and enhanced in the presence of inhibitor of mitochondrial respiration, suggesting that these pathways reduce and oxidise, respectively, the plastoquinone pool in the dark in this alga. Nigericin, which prevents the formation of proton gradient and thus qE, increased photoinhibition in *A. acetabulum* but not in *Bryopsis* sp.. Anoxia and inhibitors of the plastid terminal oxidase and mitochondrial respiration, on the other hand, enhanced photoinhibition only in *Bryopsis* sp., suggesting that oxygen-dependent pathways (including flavodiiron proteins) are important for photoprotection in the qE-deficient Bryopsidales algae.

## Introduction

Light damages the photosynthetic machinery of plants, algae and cyanobacteria and promotes production of harmful reactive oxygen species (ROS). Photosystem II (PSII) is a major target of the photodamage; the rate constant of PSII photoinhibition is directly proportional to the intensity of light (Tyystjärvi & Aro 1996), but net damage only accumulates if the concurrent repair cycle fails to keep up with the rate of the damage, for example under high light (for a recent review of PSII repair, see Su et al. 2024). The repair cycle is also sensitive to ROS (Nishiyama et al. 2001; Toriu et al. 2023). Multiple mechanisms have been suggested to lead to PSII damage: direct absorption of radiation (mostly UV and blue light) by the manganese ions of the oxygen evolving complex (Hakala et al. 2005; Ohnishi et al. 2005), oxidation by singlet oxygen (Vass 2012; Mattila et al. 2022) and oxidation by the long-lived PSII reaction centre radicals P680^+^ or TyrZ^•^ (Jegerschöld et al. 1990; Mattila et al. 2022). Photosystem I (PSI) photoinhibition, on the other hand, occurs if the electron flow from PSII exceeds the capacity of PSI electron acceptors, for example, under fluctuating light (Suorsa et al. 2012), which leads to superoxide production and oxidative damage to PSI (Inoue 1986; Tiwari et al. 2016). Even though PSI photoinhibition occurs less frequently than PSII photoinhibition, its consequences are more severe, as no efficient PSI repair cycle exists (Larosa et al. 2018; Lima-Melo et al. 2019).

As photosynthetic organisms frequently experience excess and rapidly fluctuating light conditions, they have developed multiple photoprotective mechanisms, such as non-photochemical quenching (NPQ) of excitation energy. In the model green microalga *Chlamydomonas reinhardtii*, NPQ largely depends on the light-harvesting complex stress-related (LHCSR) proteins (Peers et al. 2009; Bonente et al. 2011; Tian et al. 2019; Perozeni et al. 2020). This type of NPQ is activated by the protonation of the thylakoid lumen and is therefore called energy-dependent quenching (qE). However, NPQ in a group of green macroalgae, Bryopsidales, is induced (and relaxed) slowly and occur independently of a proton gradient (Christa et al. 2017). The genomes of these algae appear to lack the genes for the LHCSR as well as for the Photosystem II 22 kDa (PSBS) protein (Handrich et al. 2017; Iha et al. 2021; Xu et al. 2025); the latter is the pH sensor of NPQ in plants. In plants and some green algae, low luminal pH also activates the so-called xanthophyll cycle (conversion of violaxanthin to zeaxanthin), which further enhances NPQ (Quaas et al. 2015; Girolomoni et al. 2020). While some green algae possess a functional xanthophyll cycle but lack qE (Morelli et al. 2024; Havurinne et al. 2025), in Bryopsidales, only minimal accumulation of zeaxanthin occurs during dark to light transitions (Raniello et al. 2006; Christa et al. 2017).

Traditionally, NPQ has been thought to occur solely on the PSII antenna. However, the LHCSR proteins have been observed to associate also with PSI super-complexes (Allorent et al. 2013; Kosuge et al. 2018; Mosebach et al. 2024) and in *C. reinhardtii*, the LHCSR-dependent NPQ seems to quench PSI excitation, in addition to PSII excitation (Girolomoni et al. 2019). State transitions balance excitation between PSII and PSI; reduction of the plastoquinone (PQ) pool induces phosphorylation, detachment and possibly movement of a part of PSII antenna to serve PSI (state 1 to state 2 transition; Delosme et al. 1996; Depège et al. 2003; Huang et al. 2021). The association of LHSCR with PSI has been observed to strengthen during state 2 (Allorent et al. 2013). Interestingly, Bryopsidales algae have lost, in addition to qE, the capacity for state transitions (Havurinne et al. 2025).

NPQ diminishes ROS production (Girolomoni et al. 2017; Barera et al. 2021; Troiano et al. 2021) as well as photoinhibition of PSII (Jahns et al. 2000; Tyystjärvi 2013; Barera et al. 2021) and PSI (Roach 2020). Consequently, mutants deficient in NPQ are often sensitive to high and fluctuating light (Külheim et al. 2002; Peers et al. 2009; Cantrell & Peers 2017; Steen et al. 2022). In green algae, also state transitions are involved in photoprotection and double mutants deficient in NPQ and state transitions are shown to be more sensitive to PSII photoinhibition and produce more ROS than the respective single mutants (Allorent et al. 2013; Roach et al. 2015).

Besides NPQ and state transitions, auxiliary electron transfer pathways contribute to photoprotection. Plastid terminal oxidase (PTOX) oxidizes the PQ pool by reducing oxygen (for a review, see Nawrocki et al. 2015). Under constant light, PTOX may have a minor role (Bonente et al. 2012; Peltier et al. 2024), but under fluctuating light, a *C. reinhardtii* mutant deficient in PTOX activity grows slowly, possibly due to over-reduction of the PQ pool during dark periods (Nawrocki et al. 2019). Flavodiiron proteins mediate electron transfer from PSI to oxygen (safely producing water) and are especially important for PSI photoprotection in fluctuating light (Chaux et al. 2017a; Jokel et al. 2018) but have also been shown to protect PSII (Zhang et al. 2009; Bersanini et al. 2017). Indeed, *C. reinhardtii* mutants lacking flavodiiron proteins show increased PSI-derived ROS production (Pfleger et al. 2024). In addition, mitochondrial respiration and exchange of reducing power and energy equivalents between chloroplasts and mitochondria (via malate shuttles, an ADP/ATP translocator and/or a triose phosphate transporter; Lemaire et al. 1988; for a review, see Burlacot et al. 2019) are important (Peltier et al. 2024), especially in high light (Kaye et al. 2019; Huang et al. 2023). In *C. reinhardtii*, state transitions are essential for growth when mitochondrial respiration is defective (Cardol et al. 2009). Finally, cyclic electron transport from PSI to the PQ pool via the type II NADPH dehydrogenase (NDA2) and/or the proton gradient regulation 5 (PGR5)/PGR5-like (PGRL1) routes produces extra ATP, possibly powering PSII repair (Huang et al. 2018) and protecting PSI in fluctuating light (Jans et al. 2008; Chaux et al. 2017b; Jokel et al. 2018; Nikkanen et al. 2020). It has been suggested that the PSI cyclic electron transfer is especially important under anoxic conditions, where flavodiiron proteins (and other oxygen-dependent pathways) cannot function (Tolleter et al. 2011; Alric 2014; Godaux et al. 2015).

Many Bryopsidales algae inhabit fluctuating and high light environments, such as tidal pools and yet, they lack both qE and state transitions. Therefore, we hypothesize that Bryopsidales utilize compensatory photoprotective mechanisms to survive. Here, we compared two morphologically similar green macroalgae, capable of forming large, differentiated thalli comprising a single giant tubular (siphonous) cell: the qE-deficient Bryopsidales alga *Bryopsis* sp., and a “true Ulvophyte”, the Dasycladales alga *Acetabularia acetabulum* possessing qE capacity (for a recent phylogenetic analysis of the group Ulvophyceae, see Hou et al. 2022). We show that *Bryopsis* sp. is not more vulnerable to PSII photoinhibition than *A. acetabulum*. However, the two species exhibited differing vulnerabilities to inhibitors of photoprotective pathways suggesting that in the absence of qE, Bryopsidales algae indeed rely more on other available photoprotective mechanisms.

## Materials and methods

### Algal species and culture conditions

*A. acetabulum* (strain DI1 isolated by Diedrik Menzel) and *Bryopsis* sp. (previously *B. plumosa*; KU-0990; obtained from Kobe University Macroalgal Culture Collection, Japan) were grown under laboratory conditions at 20–22 °C, in the photosynthetic photon flux density (PPFD) of 40–80 µmol m^-2^ s^-1^, under 12/12 h day/night cycle, in modified f/2 medium (prepared in 35 ppt artificial sea water; Red Sea salt, Red Sea Europe, France), as previously described (Havurinne & Tyystjärvi 2020; Cartaxana et al. 2023).

### Light treatments

Intact algal cells were illuminated for 24–50 min with white or blue light of constant or fluctuating intensity (PPFD 500–2000 µmol m^-2^ s^-1^), as indicated in the respective figure legends. Reef Pulsar SPS-8 (Tropical Marine Centre; UK) was used as the white light source and Imaging-PAM (MINI; Walz, Germany) for the blue light source (see Fig. S1A for the spectra). Light intensity was measured with a wavelength-calibrated ULM-500 Universal Light Meter with a planar sensor (Walz, Germany) in air. For oxygen measurements (see below), the white build-in LEDs of the oxygen electrode system (Oxytherm+P; Hansatech Instruments; UK) were used. For fluctuating light treatments, light was either switched on and off abruptly, or gradually increased and decreased, as indicated in the corresponding figure legends. In the latter case, light intensity increased evenly for five min and then decreased for another five min; the cycle was repeated for five times (Fig. S1B). After high light treatments (or dark controls of similar lengths), two h recovery was performed under low light (white halogen lamps; PPFD 10–20 µmol m^-2^ s^-1^). All the treatments were performed at room temperature (∼20 °C) in artificial sea water, supplemented, when indicated, with the following chemicals: 10 mM lincomycin (LM), 10 µM antimycin A (AA), 6 mM glucose, 8 U/mL glucose oxidase and 800 U/mL catalase (to induce anaerobicity), 1 mM dithiothreitol (DTT), 60 µM nigericin (NIG), 10 µM oligomycin (OM), 1 mM propyl gallate (PG) or 0.1 mM polymyxin B (PMB). Stock solutions of glucose, glucose oxidase, catalase, DTT and PMB were dissolved in distilled water, those of AA, PG and NIG in ethanol and that of OM in dimethyl sulfoxide. LM was dissolved in 20 mL of artificial sea water, where the algae were incubated overnight in darkness, as indicated. Before the treatments, algae were then removed from the LM incubation and placed in fresh artificial sea water. All other chemicals were added 20 min prior to the treatments and the algae were exposed to the chemicals throughout the treatment time.

### Chlorophyll *a* fluorescence

Pulse amplitude modulated (PAM) chlorophyll *a* fluorescence was measured with the Imaging-PAM fluorometer (Walz). Frequency of the measuring flashes (setting 2) was 1 Hz. F_V_/F_M_ (=(F_M_-F_O_)/F_M_), where F_M_ is the maximum fluorescence (during a saturating light pulse) and F_O_ minimum fluorescence of a dark-acclimated sample, was measured after at least 20 min in the dark. When photoinhibition of PSII was quantified, a decrease in the F_V_/F_M_ value in the dark in the presence of the indicated chemicals (see above) was subtracted from the final values. NPQ was quantified as (F_M_-F_M_’)/F_M_’ and PSII “operational efficiency” in light as (F_M_’-F’)/F_M_’, where F’ is fluorescence and F_M_’ is the maximum fluorescence under illumination. Relative electron transport rates (rETR) were calculated as (F_M_’-F’)/F_M_’ x 0.5 x 0.84 x PPFD (note that the PSII light absorption cross-section was not measured from the algae and therefore only relative rates of PSII electron transfer can be calculated). During rapid light response curves, algae were illuminated with increasing intensities of blue light (PPFDs 4, 17, 39, 57, 84, 138, 178, 240, 309, 373, 448, 540 and 645 µmol m^-2^ s^-1^), for 60 s each, after which a saturating pulse was fired to calculate rETR and NPQ. To obtain values of alpha (initial slope of the light response curve), maximum rate of relative electron transfer (rETR_MAX_) and minimum saturating irradiance (I_K_), the curves were fitted to the model of Eilers and Peeters (1988) in Microsoft Excel.

### Isolation of thylakoid membranes

Thylakoid membranes of *Bryopsis* sp. were isolated, after breaking the cells by grinding in ice-cold mortar, according to Hakala et al. (2005). Isolations were conducted before and immediately after the light treatments (see above) and maximum oxygen evolution activity of PSII (see below) was measured immediately after the isolations. Chlorophyll contents of the thylakoid samples were quantified with a spectrophotometer according to Porra et al. (1989), after ∼1 h extraction in acetone (in the dark).

### Oxygen measurements

Oxygen concentrations were measured with oxygen electrode (Oxytherm+P; Hansatech Instruments) at 20 °C in a 2 mL closed chamber (except in the case of measurements of oxygen concentration in the presence of glucose, glucose oxidase and catalase, where the chamber was left open, to mimic the photoinhibition experiments). Intact algae were incubated in darkness, in constant white high light (PPFD 500 µmol m^-2^ s^-1^) or in low white light (PPFD ∼20 µmol m^-2^ s^-1^), as indicated, in artificial sea water. After the illumination, algal samples were gently dried with a tissue paper and weighed with an analytical scale; the oxygen production/consumption was calculated on the basis of the fresh weight. Isolated thylakoids were incubated in a buffer containing 40 mM Hepes-KOH (pH 7.4), 330 mM sorbitol, 1 M betaine monohydrate, 5 mM MgCl_2_, 5 mM NaCl, 1 mM KH_2_PO_4_, 5 mM NH_4_Cl, 0.5 mM 2,6-dichloro-1,4-benzoquinone (DCBQ) and 0.5 mM Potassium hexacyanoferrate(III). Maximum (light-saturated) PSII oxygen evolution activity of thylakoids was measured under saturating light (PPFD 4000 µmol m^-2^ s^-1^). The rates were calculated using a linear regression at 40–90 s after switching on the light; the rate of oxygen consumption under dark prior to switching on the saturating light was subtracted from final results. The rates were quantified based on chlorophyll content of the thylakoid sample.

### Figures and statistical analyses

Figures were prepared with the ggplot2 v3.4.1 package of R (R Core Team 2021; Wickham 2016). T-tests (heteroscedastic) were calculated in Microsoft Excel, using a Bonferroni correction.

## Results

### *Bryopsis* sp. displayed slow PSII repair

To test if the absence of qE makes Bryopsidales algae susceptible to PSII photoinhibition, *Bryopsis* sp. and a related alga with qE capacity (*A. acetabulum*) were illuminated with white high light of either constant (PPFD of 500 µmol m^-2^ s^-1^) or gradually fluctuating (PPFD of 0 to 1000 µmol m^-2^ s^-1^) intensity (Fig. 1A); the light dose given was equal in both treatments (see Material and methods and Fig. S1 for details of the light treatments). After the high light treatment, algae were allowed to recover for two h under low light (PPFD of 10–20 µmol m^-2^ s^-1^; Fig. 1B). The treatments were also conducted in the presence of several inhibitors of NPQ, cyclic, pseudo-cyclic and mitochondrial pathways, as well as with lincomycin (Fig. 1C), which prevents the repair of damaged PSII units. PSII photoinhibition was quantified with the chlorophyll *a* fluorescence parameter F_V_/F_M_ (after 20 min in the dark). Due to slightly lower F_V_/F_M_ values in *Bryopsis* sp. than in *A. acetabulum*, already before the treatments (Fig. S2), and because several of the used inhibitors affected the F_V_/F_M_ values in the dark (Fig. S3), all results were normalized to their respective initial F_V_/F_M_ values (prior to the high light treatments but after addition of an inhibitor). In addition, a decrease in the F_V_/F_M_ value during a 50 min dark incubation (corresponding to the duration of the high light treatment) in the presence of the chemicals (Fig. S3) was taken into account.

**Fig. 1.**
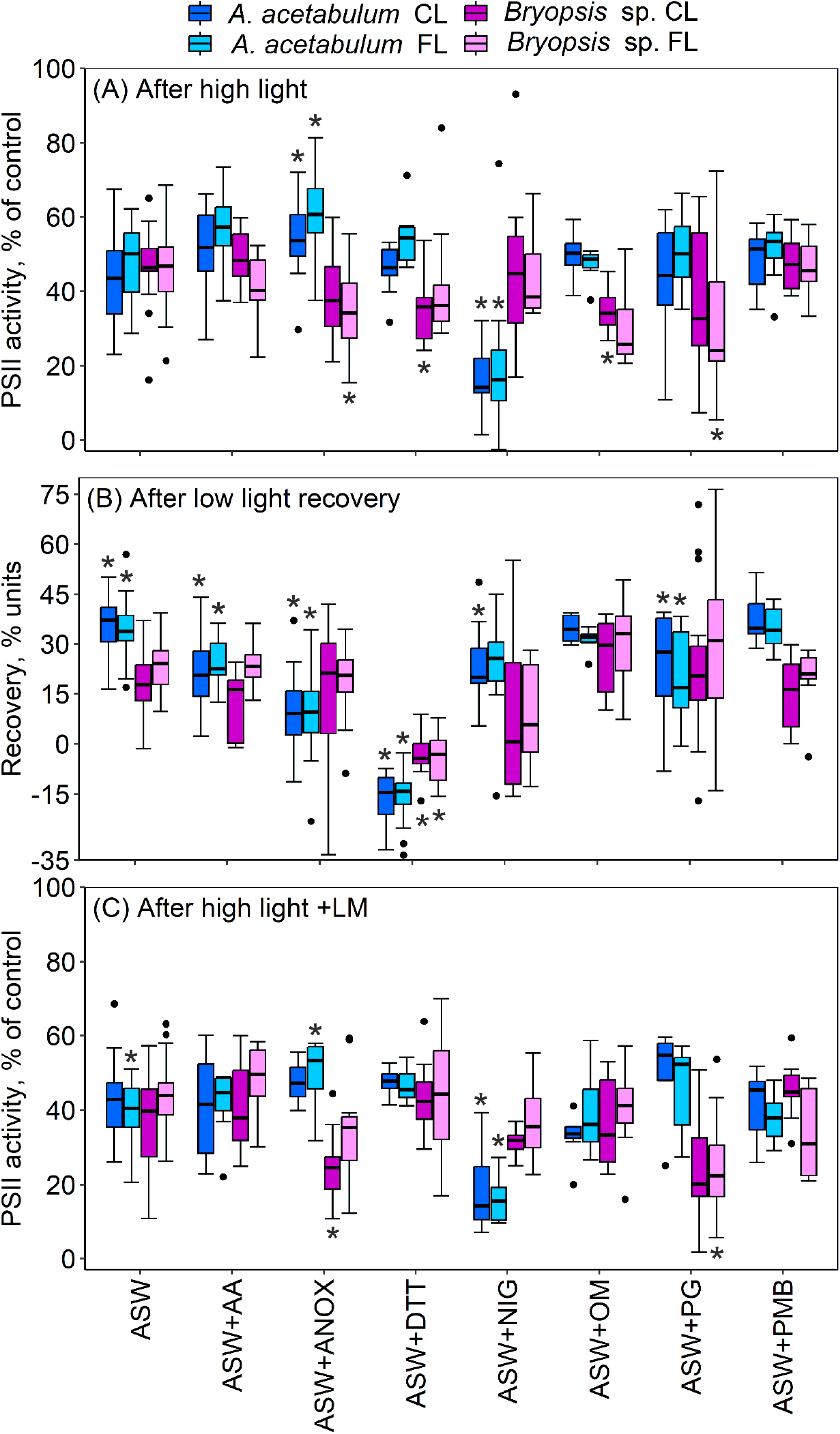
PSII photoinhibition and recovery in *Acetabularia acetabulum* and *Bryopsis* sp.. The algae were illuminated for 50 min with constant (CL; white light of PPFD 500 µmol m^-2^ s^-1^) or gradually fluctuating (FL; white light of PPFD 0 to 1000 µmol m^-2^ s^-1^; Fig. S1) light (A), after which they were let to recover at low light (PPFD 10–20 µmol m^-2^ s^-1^) for two h (B). The cumulative amount of light was equal in both the treatments. The high light treatment was also conducted with algae pre-incubated overnight in darkness in the presence of lincomycin (LM; C). All treatments were conducted at room temperature in artificial seawater (ASW) supplemented with the indicated chemicals: AA (antimycin A), ANOX (glucose, glucose oxidase and catalase; to induce anaerobicity; Fig. S4), DTT (dithiothreitol), NIG (nigericin), OM (oligomycin), PG (propyl gallate) or PMB (polymyxin B). PSII activity was assayed with the chlorophyll *a* fluorescence parameter F_V_/F_M_, measured after 20 min in the dark, and quantified as the percentage of the initial (control) F_V_/F_M_ value (after addition of the chemicals but prior to the illumination). A decrease in the F_V_/F_M_ value during a 50-min dark incubation in the presence of the chemicals (Fig. S3) was taken into account. Recovery was quantified as the difference in the percentage units between high light and recovery ((F_V_/F_M_ recovery - F_V_/F_M_ high light)/(F_V_/F_M_ initial) x 100). The box plots show medians, 2^nd^ and 3^rd^ quartiles, error bars show minimum and maximum values and dots show outliers (> 1.5 times the interquartile range), calculated based on 6–28 biological replicates. The asterisks highlight chemical treatments that statistically significantly differ from their respective treatments in ASW alone, or in the case of ASW in (B), between *A. acetabulum* and *Bryopsis* sp., or in the case of ASW in (C), between LM-treated and non-treated samples (see Tables S1–4 for details).

There were no statistically significant differences in the amount of PSII photoinhibition (decrease in the F_V_/F_M_ values due to the high light treatment) between the two algae, nor between the constant and fluctuating light treatments (Fig. 1A; tested for the samples illuminated in artificial sea water without inhibitors; Tables S1–4). The small increase in photoinhibition in the lincomycin-treated samples (statistically significant only in fluctuating light in *A. acetabulum*), compared to the samples illuminated in the absence of lincomycin, suggest that little repair occurred during the 50-min time course of the high light treatment (cf. Fig. 1A and C). On the other hand, F_V_/F_M_ values clearly increased during the two-h recovery period (without lincomycin) after the high light treatments (Fig. 1B). Without inhibitors, the F_V_/F_M_ values reached 89 ± 7.6 % (after constant light) or 83 ± 6.2 % (after fluctuating light) of the initial values in *A. acetabulum* and 65 ± 7.5 % (after constant light) or 69 ± 7.7 % (after fluctuating light) in *Bryopsis* sp. (without inhibitors); the amount of recovery was statistically significantly lower in *Bryopsis* sp. than in *A. acetabulum* (Fig. 1B; Tables S1–4).

To test whether the slow recovery in the F_V_/F_M_ parameter values in *Bryopsis* sp. really reflected poor PSII repair and not high amounts of sustained (i.e., slowly relaxing) NPQ, thylakoid membranes were isolated from the *Bryopsis* sp. cells, before and after the fluctuating high light treatment and in addition, after the low light recovery period. PSII activity was then quantified by measuring maximum oxygen evolution capacity of the isolated thylakoids. The results confirm that the PSII repair occurred slowly in *Bryopsis* sp. (Fig. 3). Photoinhibition was observed to occur slightly slower in the oxygen evolution data, compared to the fluorescence data, which may derive from the fact that the high concentration of algae used (necessary to obtain enough material for thylakoid isolations) may have caused shading or from the relatively high variation in the control oxygen evolution activity (Fig. 3).

To get more insights into the repair reactions of the two algae, the amount of recovery in each sample (without inhibitors) was plotted against the amount of photodamage in the same sample (Fig. 2). Interestingly, the more photodamage was observed in an individual algal sample, the higher was the rate of the recovery, in both algae (Fig. 2).

**Fig. 2.**
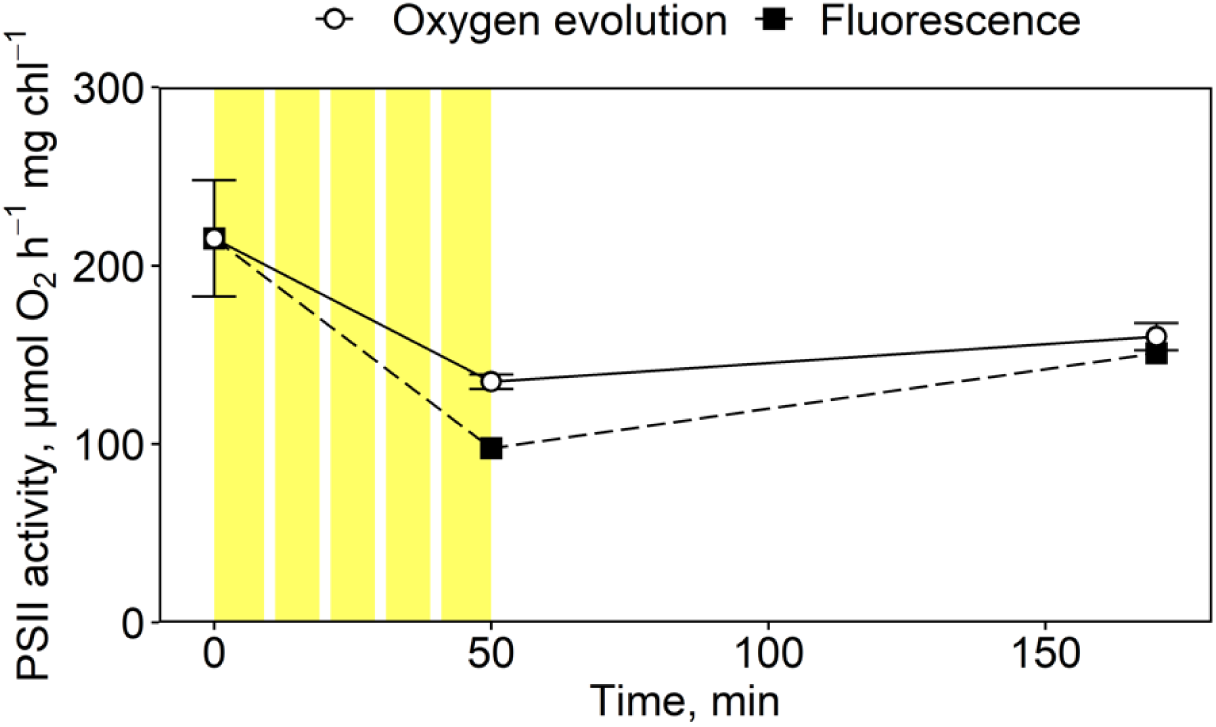
PSII photoinhibition and recovery in *Bryopsis* sp., assayed with PSII oxygen evolution (open circles and the solid line), compared to a similar experiment assayed with a chlorophyll *a* fluorescence method (black squares and the dashed line; Fig. 1). The algae were illuminated for 50 min with gradually fluctuating white light (illustrated with the yellow panels; PPFD of 0 to 1000 µmol m^-2^ s^-1^; Fig. S1) and let to recover for two h at low light (PPFD 10–20 µmol m^-2^ s^-1^). All treatments were conducted at room temperature in artificial seawater. Thylakoid membranes were isolated before and after the high light treatment and after the recovery period. Immediately after the isolations, maximum rates of oxygen evolution by PSII in the thylakoids (open circles) were measured under saturating light (PPFD 4000 µmol m^-2^ s^-1^) and in the presence of artificial electron acceptors, and quantified based on chlorophyll (chl) content of the sample. The symbols show averages and error bars standard deviations calculated based on three biological replicates (separate isolations). PSII activity estimated with the F_V_/F_M_ chlorophyll *a* fluorescence parameter (for the original data, see Figs 1A and 1B) was normalised to the control (starting) value of the oxygen evolution data, to facilitate comparison.

**Fig. 3.**
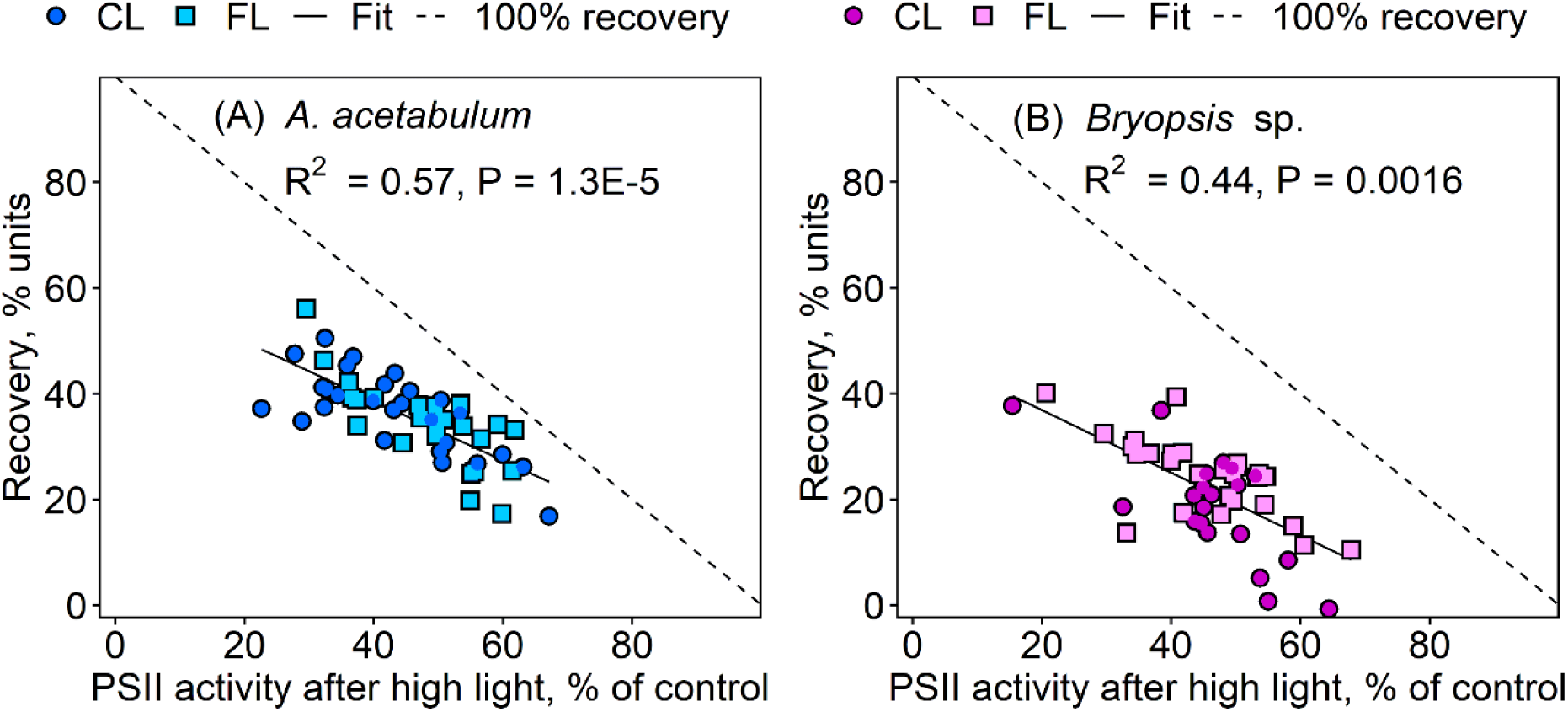
Correlations between PSII photoinhibition (PSII activity after high light, % of control) and recovery in *Acetabularia acetabulum* (A) and *Bryopsis* sp. (B). The algae were illuminated for 50 min with constant (circles; CL; white light of PPFD 500 µmol m^-2^ s^-1^) or gradually fluctuating (squares; FL; white light of PPFD 0 to 1000 µmol m^-2^ s^-1^; Fig. S1) light, after which they were let to recover at low light (PPFD 10–20 µmol m^-2^ s^-1^) for two h. PSII activity was quantified with the chlorophyll *a* fluorescence parameter F_V_/F_M_, after 20 min in the dark. All treatments were conducted at room temperature in artificial seawater. For the averaged data, see Fig. 1. The solid lines show best fits to linear equations (statistics are highlighted in the corresponding panels) and the dashed lines represent (theoretical) full recovery. The symbols represent individual biological replicates.

### Inhibitors of NPQ formation and auxiliary electron transfer pathways affected *A. acetabulum* and *Bryopsis* sp. differently

To reveal the potentially differing importances of several photoprotective pathways for *A. acetabulum* and *Bryopsis* sp., the above-described high light treatments were conducted in the presence of antimycin A (AA; an inhibitor of the PGR5/PGRL1-mediated cyclic electron transfer route), under anoxia (to inhibit oxygen-dependent pathways, most importantly flavodiiron activity; the oxygen concentrations remained below 25 nM during the illumination of both algae; see Fig. S4), dithiothreitol (DTT; an inhibitor of the violaxanthin to zeaxanthin conversion), nigericin (NIG; a dissipator of the proton gradient, preventing qE formation), oligomycin (OM; an inhibitor of the mitochondrial ATPase), propyl gallate (PG; an inhibitor of the PTOX) and polymyxin B (PMB; an inhibitor of the NDA2-mediated cyclic electron transfer route).

In *A. acetabulum*, anoxia diminished PSII photoinhibition in all the four high light treatments (constant and fluctuating high light, in the absence and presence of lincomycin), compared to the respective treatments in artificial seawater under ambient air; the difference was statistically significant in all the cases, except in constant light in the presence of lincomycin (Fig. 1A and B; Tables S1 and S2). In *Bryopsis* sp., however, anoxia enhanced photoinhibition (the effect was consistent in all the treatments but statistically significant in the case of fluctuating light without lincomycin and in constant light with lincomycin; Tables S3 and S4). In *A. acetabulum*, NIG statistically significantly enhanced photoinhibition in all the high light treatments, compared to the treatments without NIG, but had little effect on photoinhibition in *Bryopsis* sp.. On the other hand, PG increased photoinhibition in *Bryopsis* sp. in all the high light treatments (statistically significantly in the case of fluctuating light, both in the absence and presence of lincomycin) while no statistically significant differences were found in *A. acetabulum*. In *Bryopsis* sp., DTT and OM increased photoinhibition (statistically significantly in constant light), but the effect was seen only in the absence of lincomycin (Fig. 1A and B).

In *A. acetabulum*, recovery (increase in the F_V_/F_M_ values during two h under low light after the high light treatment) was negatively affected by the presence of AA, DTT, NIG, PG and anoxia (Fig. 1B; Tables S1 and S2). In *Bryopsis* sp., on the other hand, only the presence of DTT statistically significantly decreased recovery (Fig. 1B; Tables S3 and S4). Interpretation of the recovery data is, however, complicated by the fact that especially AA, DTT, NIG and PG decreased PSII activity also in samples not previously treated with high light (Fig. S3) and, therefore, their negative effects on the recovery could reflect inhibition of PSII activity in the dark rather than inhibition of the repair reactions specifically.

### During fluctuating light, fluorescence kinetics greatly differed between *A. acetabulum* and *Bryopsis* sp

To better understand how *A. acetabulum* and *Bryopsis* sp. behave under constant and fluctuating high light, chlorophyll *a* fluorescence kinetics were recorded under blue light (Fig. 4). Both during constant and fluctuating light, fluorescence initially increased, upon switching the light on, at a higher level in *Bryopsis* sp. than in *A. acetabulum*, and then decreased to similar, low levels, clearly below the original F_O_ level in both algae (Fig. 4A an B). Interestingly, at the end of the constant illumination and during the low light phases of the fluctuating light treatment, fluorescence behaved in a contrasting manner in the two algae; in *A. acetabulum*, fluorescence increased and in *Bryopsis* sp., it decreased (Fig. 4A and B), possibly due to relaxation of NPQ and increased photochemical quenching, respectively. During the constant high light illumination, PSII “operational yield” (F_M_’-F’)/F_M_’ remained close to zero in both algae (Fig. 4C). During the fluctuating illumination, (F_M_’-F’)/F_M_’ increased during the low light periods very similarly in both species (Fig. 4D). In *A. acetabulum*, however, the maximum values of (F_M_’-F’)/F_M_’ tended to remain constant or even slightly increase during the subsequent low light periods, while in *Bryopsis* sp., the parameter value decreased over time. While *A. acetabulum* rapidly induced NPQ upon a light exposure and rapidly relaxed a major part of it during the low light periods or darkness, NPQ induction was slow in *Bryopsis* sp., and almost no relaxation occurred during the low light periods (Fig. 4E and F). In both algae, the maximum amount of NPQ induced by the constant and fluctuating light treatments were rather similar, but in both cases, *Bryopsis* sp. induced less NPQ than *A. acetabulum*.

**Fig 4.**
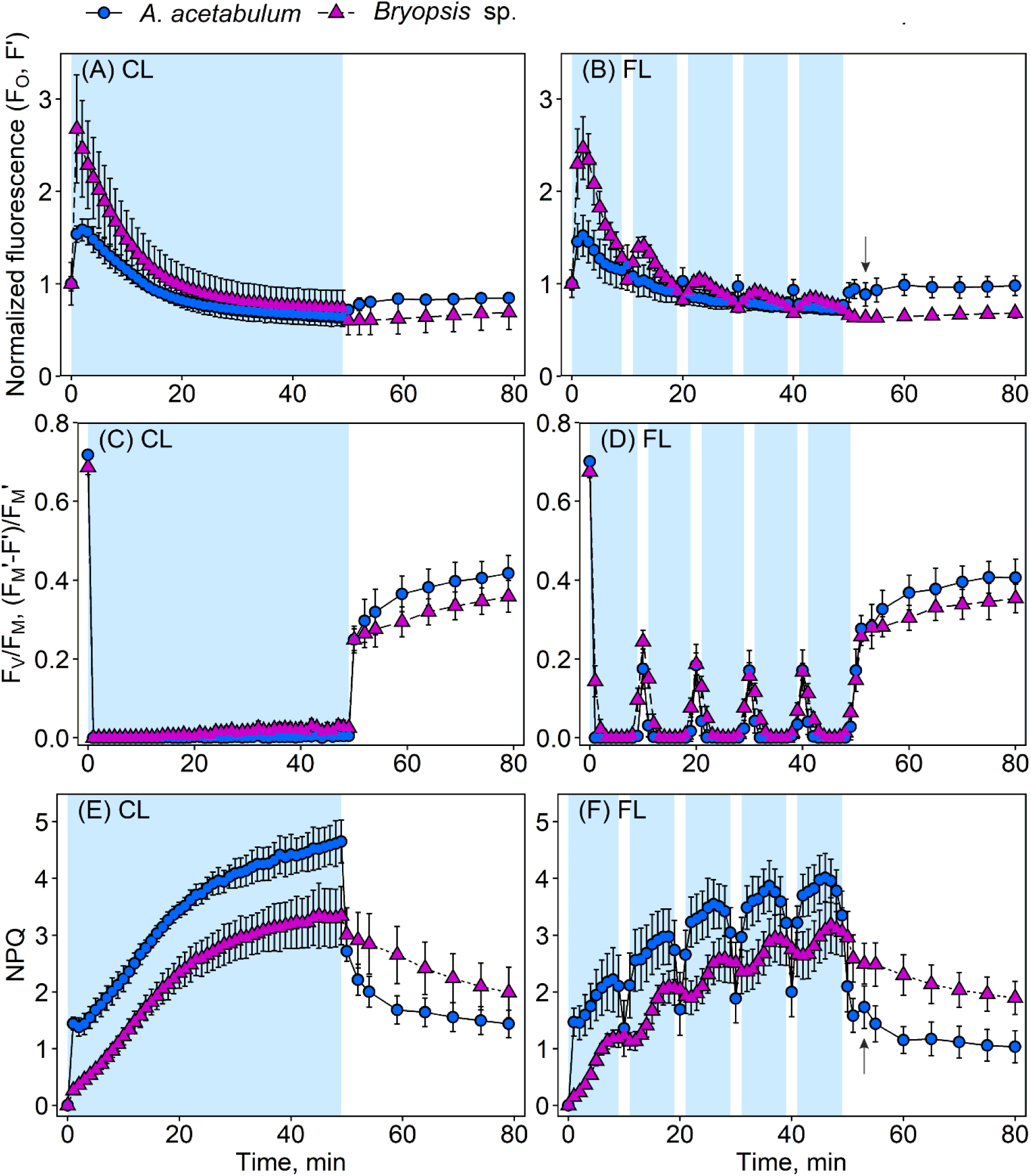
Chlorophyll *a* fluorescence under constant and fluctuating light treatments in *Acetabularia acetabulum* (circles) and *Bryopsis* sp. (triangles). The algae were dark-acclimated and fluorescence yield (F_O_, F’; normalized to the starting values; A), PSII activity (F_V_/F_M_, (F_M_’-F’)/F_M_’; B) and non-photochemical quenching (NPQ; (F_M_-F_M_’)/F_M_’; C) were measured during a 50-min treatment with constant (A, C and E; PPFD 500 µmol m^-2^ s^-1^) or gradually fluctuating (B, D and F; PPFD of 0 to 1000 µmol m^-2^ s^-1^; Fig. S1) blue light (illustrated by the blue panels) and subsequent 30-min darkness. The cumulative amount of light was equal in both the treatments. All the treatments were conducted at room temperature in artificial seawater. Symbols show averages and error bars standard deviations, calculated based on four biological replicates. The arrows in (B) and (F) highlight a post-illumination PSII activity dip and NPQ bump, respectively, in *A. acetabulum*.

### NPQ bump and PSII activity dip at the onset of darkness were affected in contrasting manner by polymyxin B and oligomycin in *A. acetabulum*

At the start of the darkness, after the fluctuating light treatment, a small dip in (F_M_’-F’)/F_M_’ and a concurring bump in NPQ was observed after the initial increase and decrease, respectively, in *A. acetabulum* but not in *Bryopsis* sp. (Fig. 4B and F). No dip or bump were observed after the constant light treatment, and therefore, to test if the strong NPQ at the end of the constant light treatment prevented the formation of the NPQ bump, the constant illumination was repeated so that the light was switched off gradually, which indeed lead to appearance of a clear NPQ bump (Fig. S5).

Next, the fluorescence measurements under fluctuating blue light were repeated under anoxia as well as in the presence of AA, DTT, NIG, OM, PQ and PMB (Figs 5 and S6). In both algae, majority of these chemicals negatively affected (F_M_’-F’)/F_M_’; in *A. acetabulum*, the severity of the inhibition during the illumination was roughly in the order NIG > PG > AA > PMB ≈ OM ≈ DTT > anoxia (Fig. 5A). In *Bryopsis* sp., a slightly different order of effects (AA > NIG ≈ PG > DTT ≈ OM > anoxia > PMB) was observed (Fig. 5C). As expected, DTT and NIG dramatically decreased NPQ in *A. acetabulum* but had little effect in *Bryopsis* sp. (Figs 5B and 5D). Additionally, anoxia and PG slightly decreased NPQ in *A. acetabulum*. The increase in fluorescence in *A. acetabulum* during low light periods (Fig. 4B) was absent in the treatments with DTT and NIG, supporting the idea of its relation to the relaxation of NPQ, but surprisingly, also in the PG treatment (Fig. S6).

**Fig. 5.**
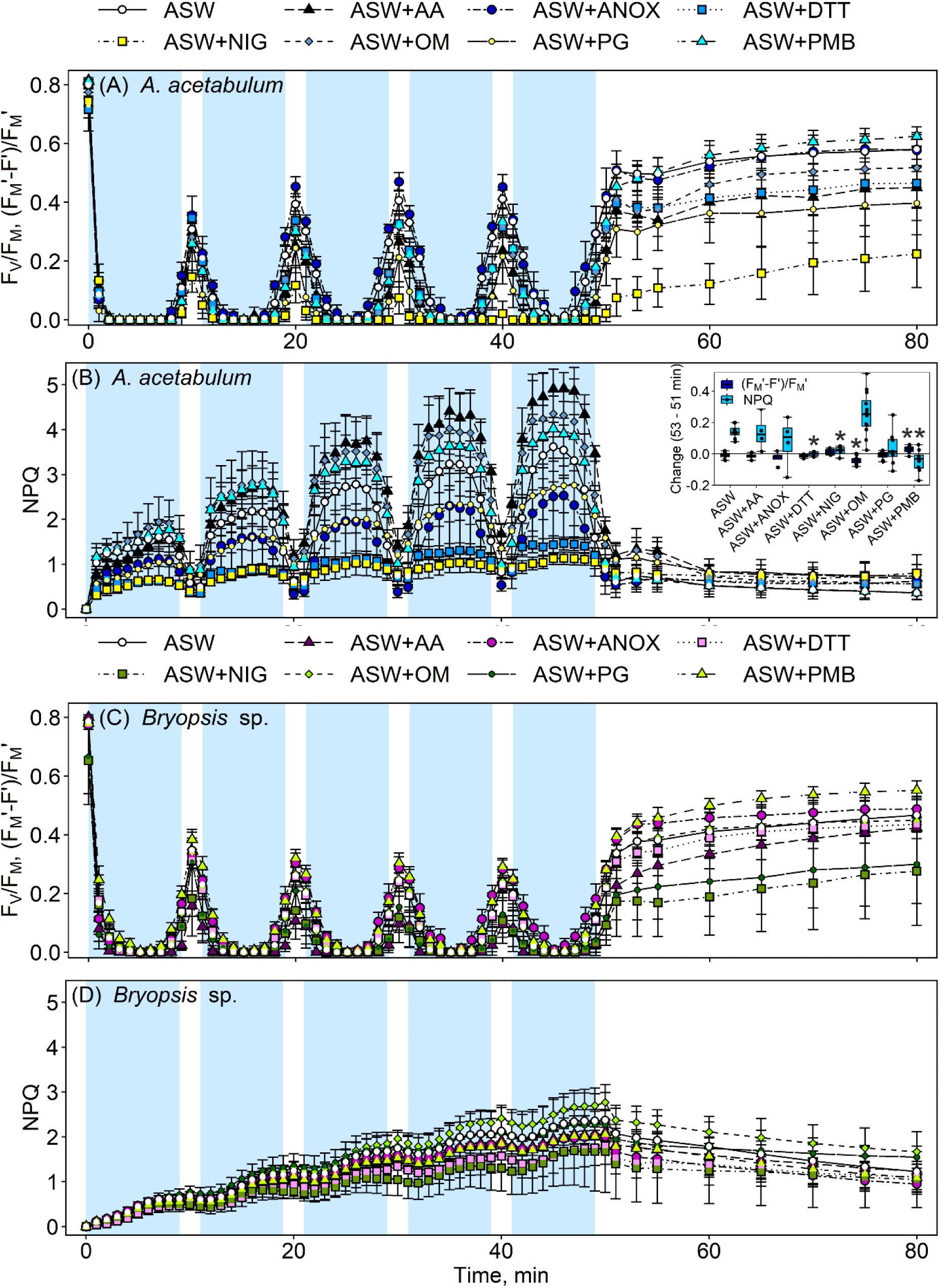
Effects of inhibitors on chlorophyll *a* fluorescence in *Acetabularia acetabulum* (A, B) and *Bryopsis* sp. (C, D). Dark-acclimated algae were illuminated for 50 min with gradually fluctuating blue light (illustrated with the blue panels; PPFD of 0 to 1000 µmol m^-2^ s^-1^; Fig. S1) and subsequently incubated for 30 min in the dark. PSII activity (A, C) was quantified as F_V_/F_M_ or (F_M_’-F’)/F_M_’ and non-photochemical quenching (NPQ; B, D) as (F_M_-F_M_’)/F_M_’. All the treatments were conducted at room temperature in artificial seawater (ASW) supplemented with the indicated chemicals: AA (Antimycin A), ANOX (glucose, glucose oxidase and catalase; to induce anaerobicity; Fig. S4), DTT (dithiothreitol), NIG (nigericin), OM (oligomycin), PG (propyl gallate) or PMB (polymyxin B). Symbols show averages and error bars standard deviations, calculated based on four to 12 biological replicates. The inset in (B) shows changes in (F_M_’-F’)/F_M_’ (dark blue boxes) and NPQ (light blue boxes) during the first minutes of darkness after the high light (quantified by subtracting the values at 51 min from the values at 53 min) in *A. acetabulum*. The horizontal line highlights zero (no change). The box plots show medians, 2^nd^ and 3^rd^ quartiles, error bars show minimum and maximum values (≤ 1.5 times the interquartile range), calculated based on four to 12 biological replicates (shown as jittered dots). The asterisks highlight treatments that significantly differ from their respective controls in ASW alone (see Table S5 for details).

In *A. acetabulum*, the NPQ bump, and a concurring tiny dip in (F_M_’-F’)/F_M_’, observed at the beginning of the darkness after the high light illuminations (Figs 4 and S5), were more pronounced in the presence of OM and disappeared in the presence of PMB (Fig 5B), though the effect of OM was statistically significant only in the case of (F_M_’-F’)/F_M_’ (Table S5). NIG and DTT also removed the NPQ peak, but the dip in (F_M_’-F’)/F_M_’ was mostly unaffected (Table S5).

### High light treatment increased electron transfer rate in *A. acetabulum* while the opposite was observed in *Bryopsis* sp

To probe the effects of NPQ and PSII electron transfer (or the capacity to keep PSII at the open state during illumination) on photoinhibition in *A. acetabulum* and *Bryopsis* sp., correlations between these parameters were calculated (Fig. S7). In both algae, (F_M_’-F’)/F_M_’ after 29 min of illumination with fluctuating light (at the PPFD ∼150 µmol m^-2^ s^-1^) and photoinhibition, estimated based on the F_V_/F_M_ values after 20 min in the dark after the illumination in the same samples, i.e. after 70 min of the experiment, showed a relatively strong correlation, especially in *Bryopsis* sp. (Figs 5 and S7A–B). NPQ after 25 min of illumination (at the PPFD 1000 µmol m^-2^ s^-1^) showed only a weak correlation with photoinhibition (Fig. S7C and D) in *A. acetabulum* and none in *Bryopsis* sp.. No correlation between (F_M_’-F’)/F_M_’ at 29 min and NPQ at the same time point was found in either of the species (Figs 5 and S7E–F).

As the openness of the PSII units under high light had a stronger relationship with the amount of PSII photoinhibition in *Bryopsis* sp. than in *A. acetabulum* (Fig. S7A and B), electron transfer was further studied by measuring rapid light response curves with chlorophyll *a* fluorescence (under blue light; Fig. S8) from the two algae before and after the white high light (constant and fluctuating) treatments. To obtain alpha (the initial slope of the rapid light response curve), rETR_MAX_ (the maximum rate of the relative electron transfer) and I_K_ (the minimum saturating irradiance) values, the light response curves were fitted to the model of Eilers and Peeters (1988). Unsurprisingly, the alpha parameter decreased after the high light treatments in both algae (Fig. 6A and B). rETR_MAX_ and I_K_, however, increased in *A. acetabulum* (though the increase was statistically significant only in the case of I_K_ after fluctuating light; Fig. 6, Table S6). In *Bryopsis* sp., on the other hand, rETR_MAX_ slightly decreased after high light (statistically significantly after fluctuating light; Fig. 6, Table S7).

**Fig. 6.**
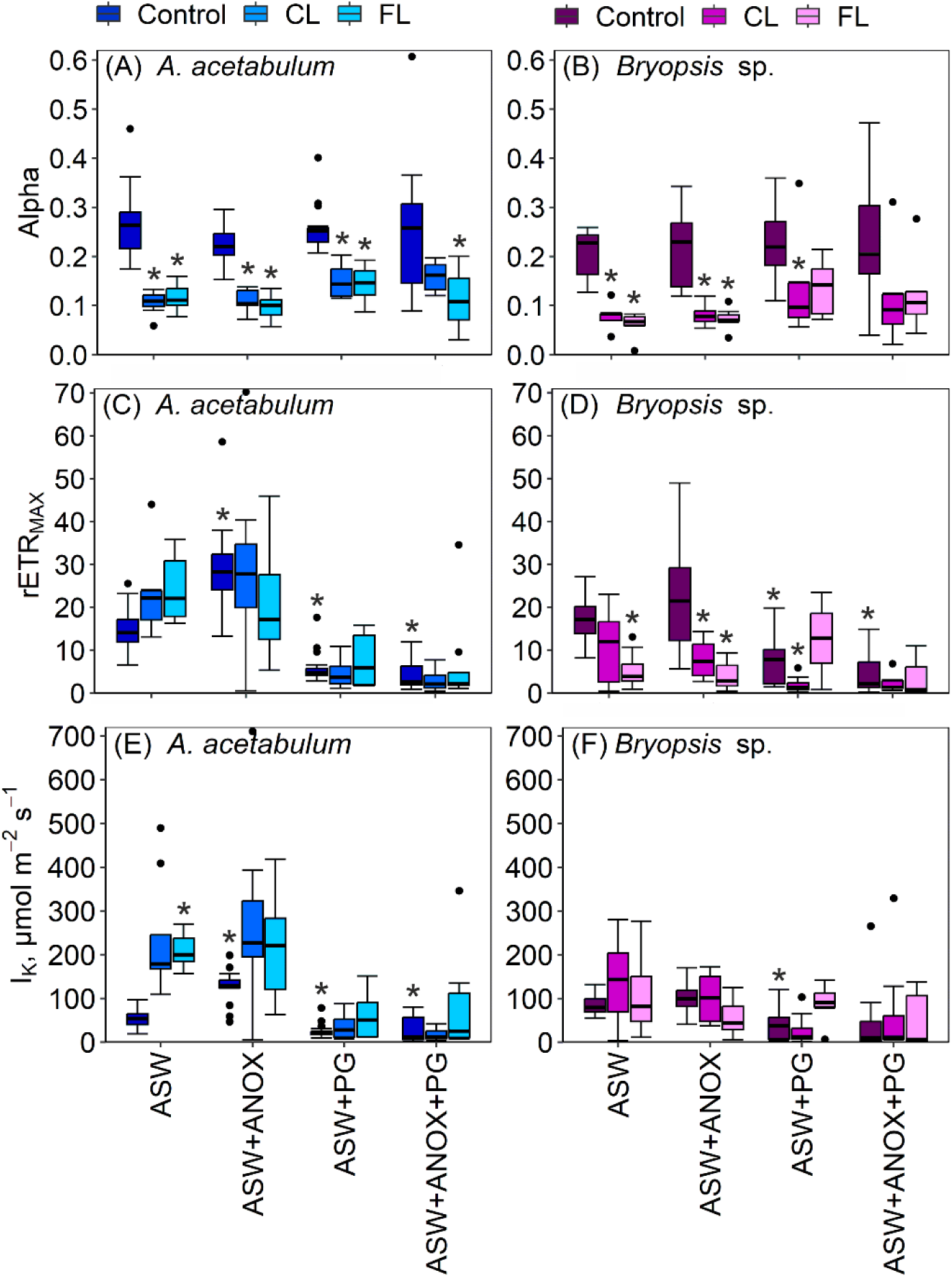
Parameters of rapid light response curves, measured with chlorophyll *a* fluorescence before (Control) and after high light treatments (constant and fluctuating) from *Acetabularia acetabulum* (A, C, D) and *Bryopsis* sp. (B, D, F). The algae were given a 50-min treatment with constant (CL; PPFD 500 µmol m^-2^ s^-1^) or gradually fluctuating (FL; PPFD of 0 to 1000 µmol m^-2^ s^-1^; Fig. S1) white light. The cumulative amount of light was equal in both the treatments. Light response curves were measured after subsequent 20-min acclimation in low light (PPFD 10–20 µmol m^-2^ s^-1^). During the light response curves, the algae were illuminated for 60 s with increasing intensities of blue light, after which a saturating pulse was fired to calculate relative rates of electron transfer. Light curves were measured at room temperature in artificial sea water (ASW), supplemented with the following chemicals (added 20 min prior to the measurements): glucose, glucose oxidase and catalase (ANOX; to induce anaerobicity; Fig. S4), propyl gallate (PG) or both, as indicated. To calculate alpha (initial slope of the light response curve; A–B), maximum electron transfer rate (rETR_MAX_; C–D) and minimum saturating intensities (I_K_; E–F), the curves were fitted to the model of Eilers and Peeters (1988). See Fig. S8 for the original fluorescence curves and examples of the fits. The box plots show medians, 2^nd^ and 3^rd^ quartiles, error bars show minimum and maximum values and dots show outliers (> 1.5 times the interquartile range), calculated based on eight to 16 biological replicates. The asterisks highlight treatments that statistically significantly differ from their respective controls, or in the case of control, from the chemical-treated controls (see Tables S6 and S7 for details).

To confirm the different responses of electron transfer to high light treatments in the two algae, rates of oxygen evolution (and consumption) by intact cells were measured (Fig. 7). In accordance with the fluorescence measurements, the rate of oxygen evolution increased up to ∼25 min under constant white high light in *A. acetabulum*, whereas *in Bryopsis* sp., high rates of oxygen evolution were observed almost immediately after switching on the light, after which the rate steadily decreased (Fig. 7).

**Fig. 7.**
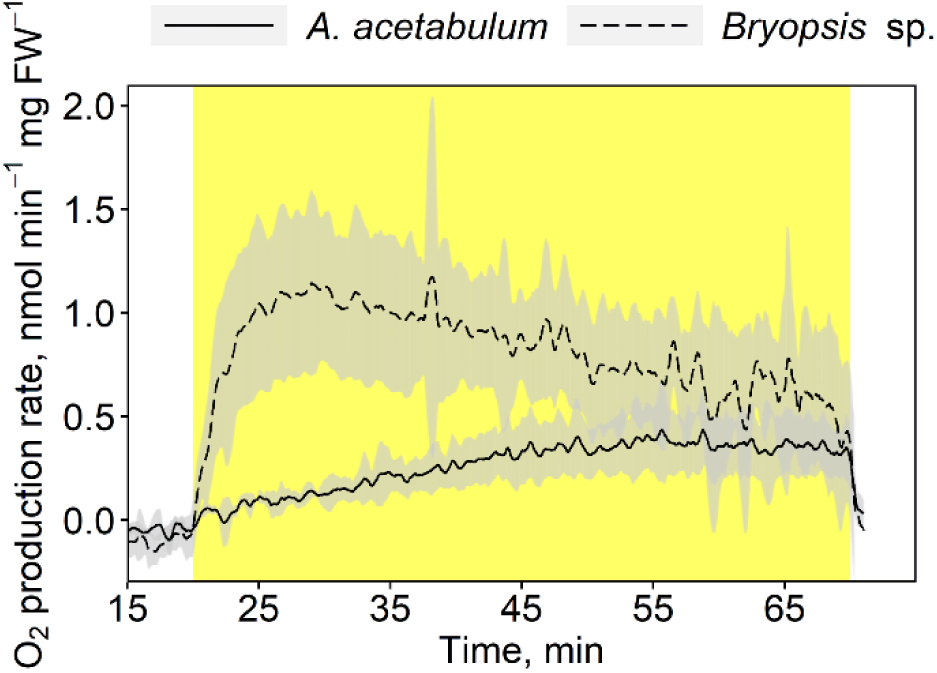
Net consumption (in the dark) and evolution (in the light) of oxygen by intact cells of *Acetabularia acetabulum* and *Bryopsis* sp. during a 20-min darkness and subsequent 50-min illumination with constant white light (illustrated with the yellow panel; PPFD 500 µmol m^-2^ s^-1^) at 20 °C in artificial sea water. The oxygen production/consumption has been calculated on the basis of the fresh weight (FW) of the sample. The lines show averages and the shaded areas standard deviations calculated based on five to six biological replicates.

Anoxia and PG had opposite effects on photoinhibition in *A. acetabulum* and in *Bryopsis* sp. (Figs 1A and C). Therefore, their effects on electron transfer were also studied by adding the chemicals 20 min prior to the measurements of the light response curves. Anoxia increased rETR_MAX_ and I_K_ values in non-illuminated *A. acetabulum* whereas no further increases were observed in the high light-treated samples (Fig. 6C and E). No significant effects were observed in *Bryopsis* sp. (Fig. 6D and F). PG as well as the combined treatment with anoxia and PG clearly diminished electron transfer rates in both algae (Fig. 6C–F).

NPQ formed during the light response curves was also quantified; in general, NPQ induction decreased after the high light treatments, especially in *Bryopsis* sp. (Fig. 8). In addition, in *A. acetabulum*, a transient peak in NPQ appeared after the high light treatments, which did not form in the presence of PG (Fig. 8E and G). In the PG-treated *A. acetabulum* samples, high light exposed algae also induced more NPQ than in the absence of PG, which may be related to the slow rates of electron transfer in these samples. In accordance with the previous fluorescence measurements (Fig. 5D), the chemicals had little effect on NPQ in *Bryopsis* sp. (Fig. 8).

**Fig. 8.**
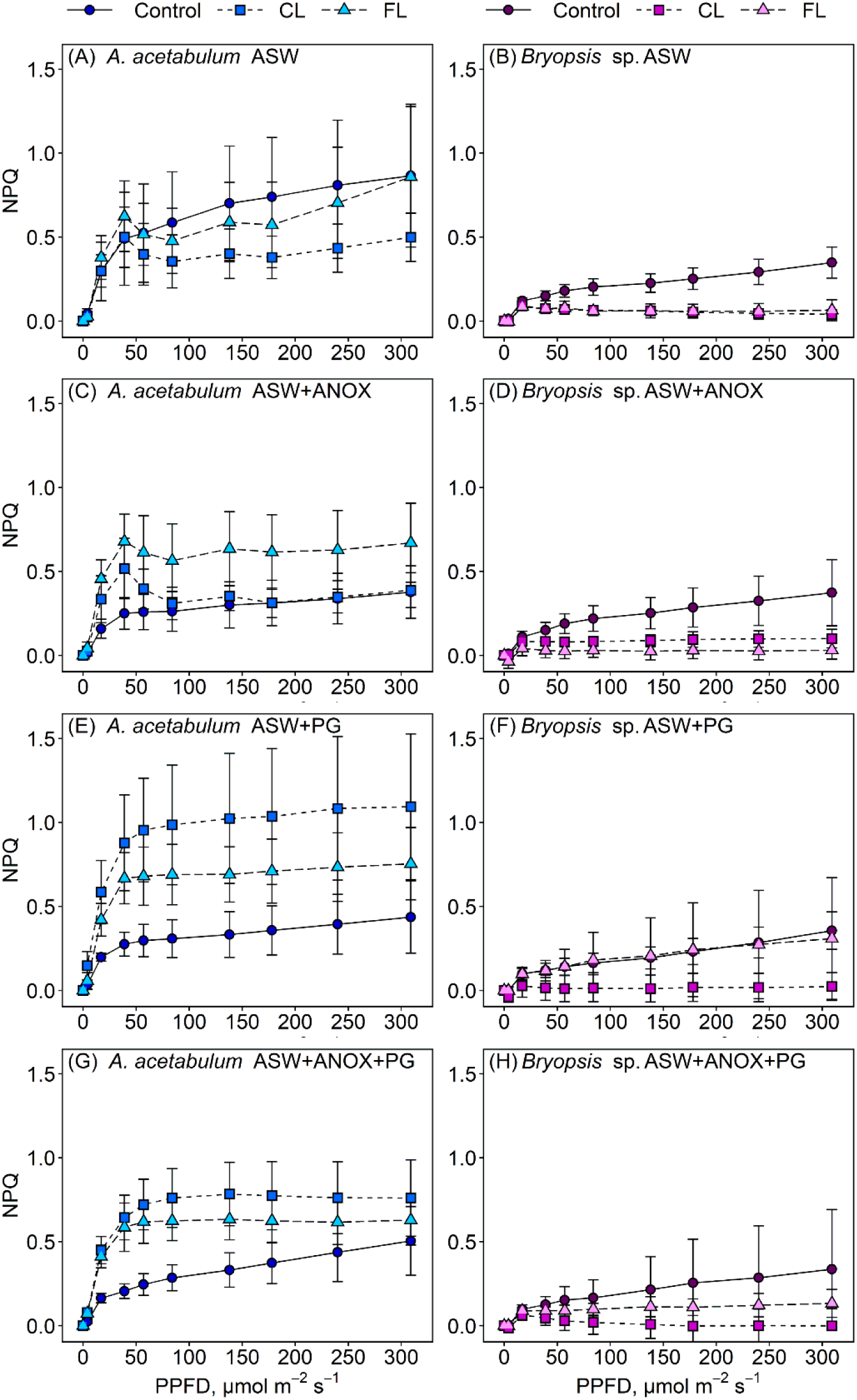
Non-photochemical quenching (NPQ; (F_M_-F_M_’)/F_M_’), measured with chlorophyll *a* fluorescence, during rapid light response curves before (Control) and after high light (constant or fluctuating) treatments in *Acetabularia acetabulum* (A, C, E, G) and *Bryopsis* sp. (B, D, F, H). The algae were given a 50-min treatment with constant (CL; PPFD 500 µmol m^-2^ s^-1^) or gradually fluctuating (FL; PPFD of 0 to 1000 µmol m^-2^ s^-1^; Fig. S1) white light. The cumulative amount of light was equal in both the treatments. Light response curves were measured after subsequent 20 min acclimation in low light (PPFD of 10–20 µmol m^-2^ s^-1^). During the light response curves, the algae were illuminated for 60 s with increasing intensities of blue light, as indicated, after which a saturating pulse was fired to calculate NPQ. Light response curves were measured at room temperature in artificial seawater (ASW) (A and B), supplemented with the following chemicals (added 20 min prior to the measurements): glucose, glucose oxidase and catalase (ANOX; to induce anaerobicity; Fig S4; C and D), propyl gallate (PG; G and F) or both (E and F), as indicated. The symbols show averages and error bars standard deviations calculated based on eight to 16 biological replicates.

### The frequency of light fluctuations did not significantly affect photoinhibition in either of the algae

The fluctuating light used here so far was relatively mild as the light intensity gradually increased from zero to the PPFD 1000 µmol m^-2^ s^-1^ during a time-course of five min (see Fig. S1). To test if *A. acetabulum* and *Bryopsis* sp. can tolerate also faster fluctuations, additional illumination treatments were conducted where the light intensity was immediately increased from zero to the PPFD 2000 µmol m^-2^ s^-1^. Furthermore, four different sequences of fluctuations were used; four min of high light and four min of darkness (termed very long fluctuating light; only in the case of *Bryopsis* sp.), four min of high light and one min of darkness (long fluctuating light), two min of high light and 30 s of darkness (medium fluctuating light) and one min of high light and 15 s of darkness (short fluctuating light) as well as the corresponding constant high light (Fig. 9). In *A. acetabulum*, the presence of fluctuations or their frequency did not significantly affect (F_M_’-F’)/F_M_’, PSII photoinhibition (probed by the F_V_/F_M_ parameter after 20 min of darkness after the high light) or maximum NPQ values, although the longer the darkness, the more relaxation of NPQ occurred during the dark phases (Fig. 9A and B). In *Bryopsis* sp., in contrast to *A. acetabulum*, (F_M_’-F’)/F_M_’ values during the dark periods were the lower the shorter the period was (Fig. 9). However, no clear effect on photoinhibition was observed in *Bryopsis* sp. either. In both algae, NPQ in general increased at the same rate during constant and fluctuating illuminations, which resulted higher maximum NPQ values at the end of the fluctuating light treatments, compared to the constant one, as these lasted longer (Fig. 9B and D). However, in *Bryopsis* sp., the 4-min darkness resulted in a slower build-up of NPQ, due to significant NPQ relaxation and a slow induction (as compared to *A. acetabulum*).

**Fig. 9.**
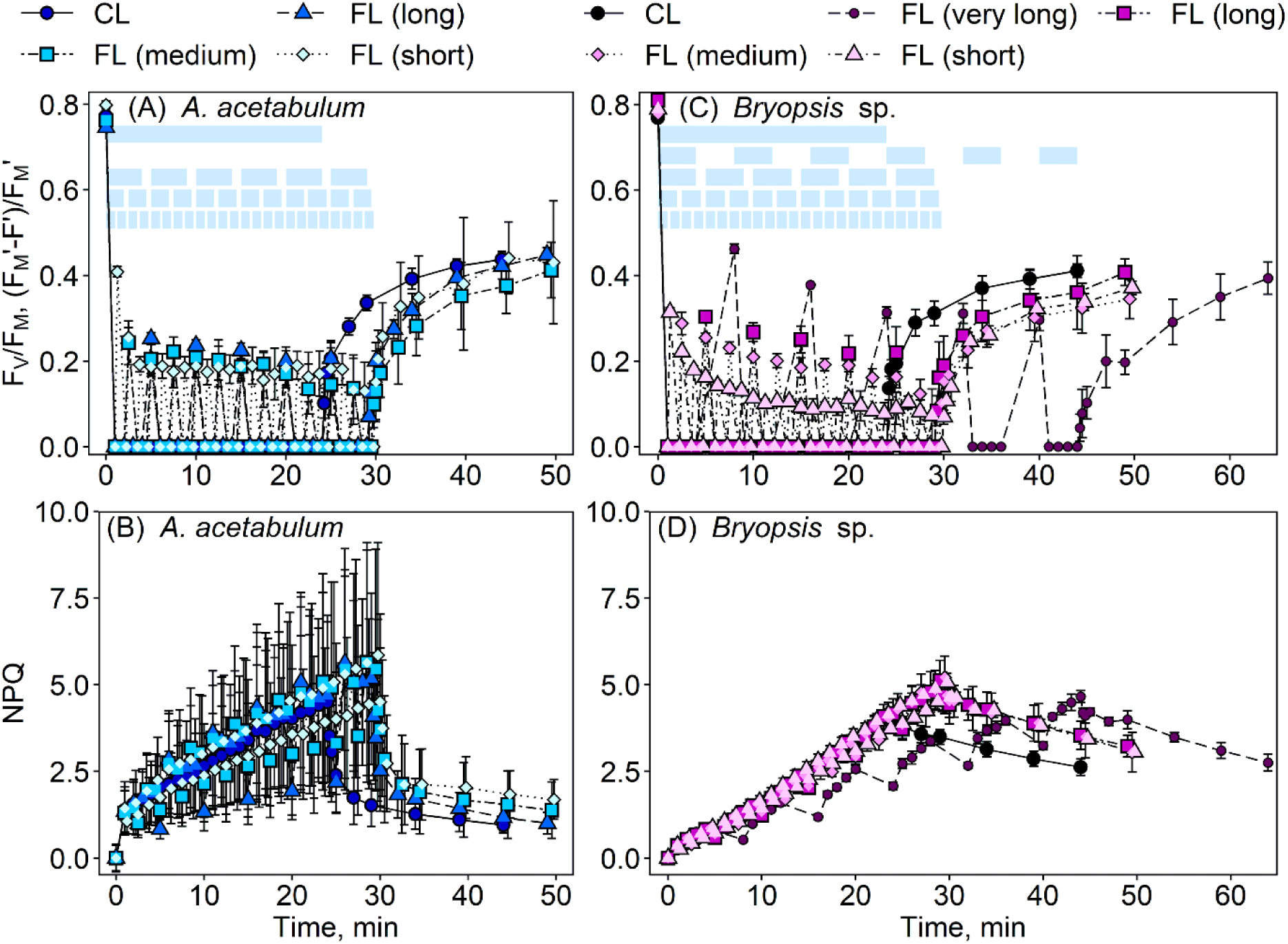
Effects of constant and different fluctuating light treatments on chlorophyll *a* fluorescence in *Acetabularia acetabulum* (A, B) and *Bryopsis* sp. (C, D). Dark-acclimated algae were given a 24-min treatment with constant (CL; PPFD 2000 µmol m^-2^ s^-1^) or 29–44-min treatments with sharply fluctuating (FL; PPFD of 0 to 2000 µmol m^-2^ s^-1^) blue light and a subsequent 20-min darkness. The fluctuating light treatments consisted of sequences of four min of high light and one min of darkness (very long FL), four min of high light and one min of darkness (long FL), two min of high light and 30 s of darkness (medium FL) and one min of high light and 15 s of darkness (short FL); the blue panels (in A, C) illustrate the different fluctuations. The cumulative amount of light was equal in all the treatments. PSII activity (A, C) was quantified as F_V_/F_M_ (after the initial 20 min of darkness) or as (F_M_’-F’)/F_M_’ (during light and the subsequent darkness) and non-photochemical quenching (NPQ) as (F_M_-F_M_’)/F_M_’ (B, D). All the treatments were conducted at room temperature in artificial seawater. The symbols show averages and error bars standard deviations, calculated based on four biological replicates.

## Discussion

At least some Bryopsidales algae are missing the fast, energy-dependent form of NPQ (qE) as well as state transitions (Christa et al. 2017; Havurinne et al. 2025), two photoprotective mechanisms previously shown to be especially important under conditions of high and fluctuating light (Külheim et al. 2002; Peers et al. 2009; Allorent et al. 2013; Roach et al. 2015; Cantrell & Peers 2017; Steen et al. 2022). Some Bryopsidales algae have indeed adapted to low light environments (Iha et al. 2021) but others inhabit shallow coastal waters, even so successfully as having become invasive and harmful (Provan et al. 2004; Klein & Verlaque 2008; Piazzi et al. 2016), and thus frequently experience extreme light fluctuations. Here, we show that the Bryopsidales alga *Bryopsis* sp. is not more susceptible to photoinhibition of PSII, under constant or fluctuating light, than *A. acetabulum* (Fig. 1), an Ulvophyceaen green macroalga with similar siphonous morphology to *Bryopsis* sp., but with clear capacities for qE (Fig. 4) and state transitions (Havurinne et al. 2025).

### *A. acetabulum* relies on NPQ while *Bryopsis* sp. utilizes oxygen-dependent pathways for photoprotection

Upon a high light (constant or fluctuating) treatment, *Bryopsis* sp. slowly induced strong NPQ that also relaxed slowly (Figs 4, 5 and 9); such a strong NPQ has been previously suggested to contribute to photoprotection in *Bryopsis corticulans* (Giovagnetti et al. 2018). Here, also *A. acetabulum* was observed to induce similar, high levels of NPQ, but as the NPQ induction occurred rapidly in *A. acetabulum*, the difference in NPQ levels between *Bryopsis* sp. and *A. acetabulum* was large at the first ∼10 min of illumination. Therefore, some damage might be expected to occur in *Bryopsis* sp. during the first minutes of high light or when the dark periods of a fluctuating light are long enough for *Bryopsis* sp. to relax substantial amount of the NPQ (Fig. 9). However, NPQ levels did not correlate well with PSII photoinhibition in either of the species (Fig. S7). In the case of *A. acetabulum*, the ability to switch NPQ rapidly on may be more important than the actual NPQ levels, as removal of the proton gradient with NIG (i.e., removal of the capacity for fast NPQ induction/relaxation) increased the rate of the photodamage to PSII while inhibition of the xanthophyll cycle with DTT (causing a clear decrease in maximum NPQ while the capacity for fast induction/relaxation remained; Fig. 5) did not (Fig. 1).

In *A. acetabulum*, anoxic (or microoxic; Fig. S4) conditions diminished PSII damage, an effect previously observed in plant leaves (Fan et al. 2016), possibly due to decreased production of singlet oxygen (Hideg et al. 1994; Pfleger et al. 2024). Interestingly, the opposite was true for *Bryopsis* sp. (Fig. 1), suggesting that oxygen-dependent pathways (flavodiiron proteins, PTOX and/or mitochondrial respiration) are more important for *Bryopsis* sp. in preventing PSII photoinhibition than for *A. acetabulum*. Alternatively, increased photoinhibition under microoxic conditions could derive from increased superoxide production by PSI, due to inactivation of the flavodiiron activity (Pfleger et al. 2024). However, usually only the ROS produced by PSII itself are considered to be relevant for PSII damage (Song et al. 2006; Kale et al. 2017). The importance of PTOX and, perhaps to a lesser degree, mitochondrial respiration for *Bryopsis* sp. is supported by the increased PSII damage in the presence of PG and OM (Fig. 1), while in *A. acetabulum*, these chemicals did not statistically significantly affect photoinhibition. However, microoxic conditions did not decrease the electron transfer rates (estimated by chlorophyll fluorescence) in either of the algae (Fig. 6), nor did they lower the PSII operational yield under fluctuating light (Fig. 5), suggesting that oxygen dependent pathways do not significantly contribute to the linear electron transport rates, but may work transiently upon sudden increases or decreases in light intensity. Indeed, transient activation of flavodiiron-dependent oxygen consumption has been observed during dark to light transitions (Santana-Sanchez et al. 2019; Saroussi et al. 2019). On the other hand, PG strongly suppressed PSII electron transfer (Figs 5 and 6). The result may not indicate high PTOX activity, however, as removal of oxygen (where no decrease in electron transfer was observed) should also hinder PTOX, but may be a side effect of PG.

PSII operational yield (i.e., (F_M_’-F’)/F_M_’) correlated with PSII photoinhibition more strongly in *Bryopsis* sp. than in *A. acetabulum* (Fig. S7). Accordingly, high light treatment lowered electron transfer rates and net oxygen evolution in *Bryopsis* sp., while in *A. acetabulum* these parameters increased (Figs 6 and 7) after high light illumination. However, the data suggest that fast photoinhibition lowers electron transfer rates in *Bryopsis* sp., rather than the low electron transfer rates causing photoinhibition, as during the early time points of the fluctuating illumination, i.e., after 1 and 9 min of illumination, weaker correlations (R^2^ = 0.48 and R^2^ = 0.51, respectively) were found between the above-mentioned parameters than in the later time points, under the same light intensities (R^2^ = 0.62; Fig. S7).

### Why does *Bryopsis* sp. repair damaged PSII units slowly?

Compared to *A. acetabulum*, *Bryopsis* sp. showed slow rates of PSII repair (Figs 1–3). Relatively high F_V_/F_M_ values (>0.55) during low tide or at noon are reported for Bryopsidales algae growing in the field (Raniello et al. 2006; Giovagnetti et al. 2018), however, complete recovery may occur also in these cases only at the evening or when the light stress has passed. Synthesis of the new D1-protein during the PSII repair cycle is known to be sensitive to ROS, due to oxidation of the elongation factors EF-G and EF-Tu (Kojima et al. 2007; Yutthanasirikul et al. 2016). Thus, the strongly suppressed repair in *A. acetabulum* under microoxic conditions (Fig. 1) could be explained by increased superoxide production by PSI (Pfleger et al. 2024). On the other hand, PSII repair was not affected by anoxia in *Bryopsis* sp. (Fig. 1). Furthermore, in both algae, repair rates increased with increasing rates of damage (Fig. 3), suggesting that the slow recovery in *Bryopsis* sp. was not due to decreased repair capacity (caused by, e.g., excess ROS production), but rather due to down-regulation of the activity of the repair cycle.

Why didn’t the algae, especially *Bryopsis* sp., repair all the damage? PSII repair requires energy (Murata & Nishiyama 2018; Yi et al. 2022) and decreased rates of protein synthesis (including the PSII reaction centre protein D1) under oxidative stress has been suggested to be a regulatory response, to prevent excessive ATP consumption, PSI damage and/or further ROS production (Tikkanen et al. 2014; Yutthanasirikul et al. 2016; Murata & Nishiyama 2018; Cheng et al. 2024). In plants, moderate photoinhibition does not decrease carbon fixation rates under high light, suggesting that plants contain “extra” PSII units (Mattila et al. 2020). Also in *A. acetabulum*, high light treatments rather increased PSII electron transfer rate (Figs 6 and 7) and thus, sustained photoinhibition may not prevent efficient photosynthesis. However, as the opposite was true for *Bryopsis* sp., the present results suggest that the algae lose opportunities for carbon fixation due to the accumulation of non-functional PSII units. In the field, photoinhibition caused no decrease in electron transfer rates in a Bryopsidales alga (Raniello et al. 2006), indicating that photosynthetic reactions are differentially regulated when the algae are grown under low light (here) and under high light (in the field). Accordingly, regulation of photosynthesis was found to be one of the upregulated gene ontology categories in high light treated *B. corticulans* (Xu et al. 2025). Alternatively, *Bryopsis* sp. may up-regulate repair rates if grown under high light, as occur in plants (e.g., Miyata et al. 2015). However, little is known about the regulation of PSII repair in (macro)algae.

ATP deficiency due to decreased mitochondrial respiration has been previously suggested to hinder repair in seagrasses under anoxia (Che et al. 2023), which also could explain low rates of repair under the microoxic conditions in *A. acetabulum*. However, *A. acetabulum* showed high electron transfer rates under anoxia (Fig. 6). In addition, inhibition of the mitochondrial respiration did not affect repair in either of the algae (Fig. 1). In plants, also PSI cyclic electron transfer is important for efficient PSII repair (Huang et al. 2018). Here, inhibition of the cyclic electron transfer did not greatly affect PSII repair in either of the algae; AA did slow down repair in *A. acetabulum*, but the effect could be due to inactivation of PSII by AA also under darkness (Fig. S3). However, cross-compensation among the different auxiliary electron pathways (Chaux et al. 2017b; Peltier et al. 2024) may also explain the lack of effect of a single inhibitor.

### The NDA2-dependent cyclic pathway is responsible for dark-reduction of the PQ pool in *A. acetabulum*

After both fluctuating and constant (if the light was switched off gradually; Fig. S5) light treatments, a small dip in the PSII operational yield and a concurrent bump in NPQ was observed in the dark in *A. acetabulum* (Figs 4 and 5), suggesting a transient reduction of the PQ pool. In green algae, dark reduction of the PQ pool can occur via both NDA2 and PGR5/PGRL1 cyclic routes (Patil et al. 2022). As the dip and the bump disappeared in the presence of PMB, while a treatment with AA had no effect (Fig. 5B), *A. acetabulum* seems to prefer the NDA2 route. A slightly bigger dip in PSII operational yield was observed also under microoxic conditions (though the difference was not statistically significant, as compared to the algae illuminated in the presence of oxygen in plain artificial sea water). Accordingly, *C. reinhardtii* has been shown to upregulate cyclic pathways under anoxia, to compensate for the lack of oxygen-dependent pathways (Alric 2014; Godaux et al. 2015).

Interestingly, both the dip in PSII operational yield and the NPQ bump were more pronounced in the presence of OM (although not statistically significantly in the case of NPQ; Fig. 5B), suggesting that in *A. acetabulum*, chloroplast to mitochondria pathways function as a sink, oxidizing the plastoquinone pool. Indeed, it has been previously shown that active mitochondrial respiration can oxidize PQ pool during light to dark transitions (Uhmeyer et al. 2017). Furthermore, mutants deficient in these pathways are sensitive to photoinhibition of both PSII and PSI, possibly due to increased ROS production and acceptor side limitation, respectively (Kaye et al. 2019; Huang et al. 2023). AA may block the mitochondrial complex III, in addition to cyclic electron transfer, but the effect of this inhibition has been shown to be small if alternative oxidase (AOX) can function (Schober et al. 2019).

We did not observe NPQ bumps or PSII activity dips in *Bryopsis* sp., however, this may be because the slow fluorescence kinetics in this alga hide the effects of the PQ pool reduction, not because dark reduction of the PQ pool does not occur in *Bryopsis* sp.. Indeed, the increased photoinhibition in the presence of OM in *Bryopsis* sp. (Fig. 1) speaks for the importance of these pathways for this alga.

### Conclusions

PSII damage under constant high or fluctuating light did not proceed faster in *Bryopsis* sp. than in *A. acetabulum*, despite the lack of rapid NPQ induction in the former alga. Previously, the protective effect of NPQ against PSII damage has been shown to be rather small (for a review, see Tyystjärvi 2013), which agrees with the present results. However, *Bryopsis* sp. did show slow PSII repair. Regardless, because the repair rates were similar after constant and fluctuating light, and actually faster in algae experiencing more damage, this result may not derive from increased damage either. To summarize, even without qE, PSII in *Bryopsis* sp. is well protected against photodamage. Different sensitivities of PSII damage, repair and electron transfer to inhibitions of NPQ formation and auxiliary pathways were observed between the two algae, suggesting that regulation of photosynthesis and photoprotection do occur differently in *A. acetabulum*, which is capable of qE, and in the qE-deficient *Bryopsis* sp.. In the absence of qE, oxygen-dependent pathways seemed to be especially important.

Even though fluctuating light can also increase PSII damage, due to increased singlet oxygen production (Davis et al. 2016), PSI is usually more sensitive to fluctuating light (e.g., Suorsa et al. 2012). The slightly more pronounced decrease in maximum electron transfer rates after fluctuating than after constant light in *Bryopsis* sp. (Fig. 6) may indeed suggest increased PSI damage. However, actual measurements of PSI activity should be conducted to test whether Bryopsidales PSI is vulnerable to fluctuating light.

## Data availability

The raw data will be available upon publication at xxx.

## Acknowledgements

This work was supported by the Finnish Cultural Foundation through the Foundation’s Post Doc Pool to H.M., by the European Research Council (ERC) under the European Union’s Horizon 2020 research and innovation program, grant agreement no. 949880 to S.C. (https://doi.org/10.3030/949880), and by Fundação para a Ciência e Tecnologia (FCT), grants 2020.03278.CEECIND to S.C. (https://doi.org/10.54499/2020.03278.CEECIND/CP1589/CT0012), CEECIND/01434/2018 to P.C. (https://doi.org/10.54499/CEECIND/01434/2018/CP1559/CT0003) and UID/50006 + LA/P/0094/2020 to CESAM (doi.org/10.54499/LA/P/0094/2020). We thank Maria Inês Silva and Diogo Marçal for technical support in the maintenance of algae.

## Supplementary figures

**Fig. S1.**
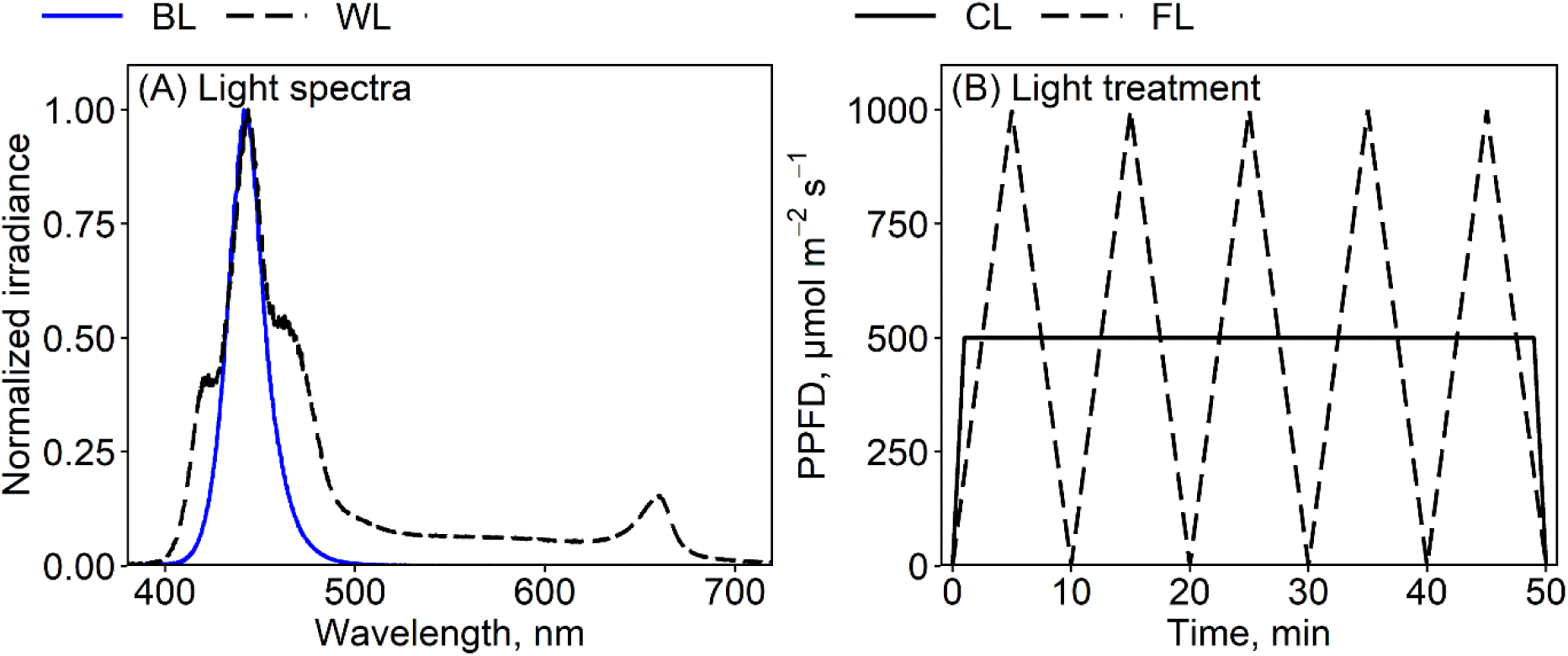
Light treatments. (A) Normalized energy spectra of the blue (BL; Imaging-PAM, Walz, Germany) and white (WL; Reef Pulsar SPS-8, Tropical Marine Centre, UK) lights used in the high light treatments in the present study. (B) A schematic presentation of the constant (CL) and gradually fluctuating (FL) light treatments. PPFD = photosynthetic (400–700 nm) photon flux density.

**Fig. S2.**
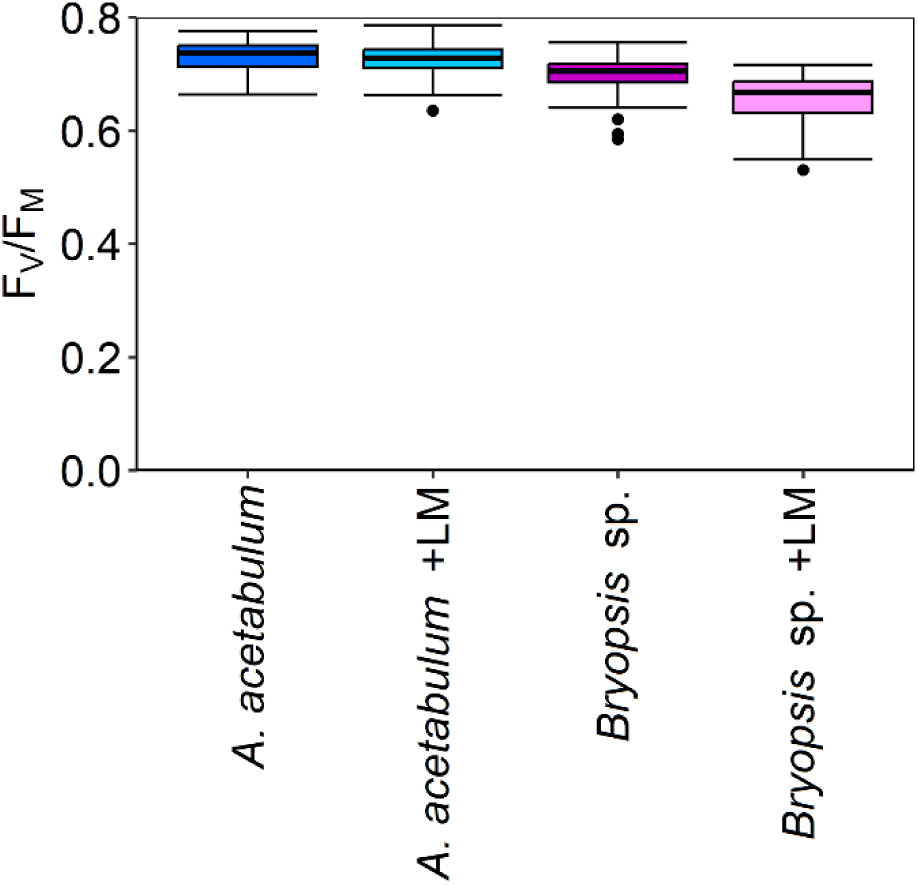
Control PSII activities of *Acetabularia acetabulum* and *Bryopsis* sp., used in the photoinhibition experiments (Fig. 1). The algae were either taken from the growth conditions or incubated overnight in the dark in the presence of lincomycin (LM), as indicated. PSII activity was quantified with the chlorophyll *a* fluorescence parameter F_V_/F_M_, after 20 min in the dark. The box plots show medians, 2^nd^ and 3^rd^ quartiles, the error bars show minimum and maximum values and dots show outliers (> 1.5 times the interquartile range), calculated based on 50 (*A. acetabulum* without LM), 34 (*A. acetabulum* with LM), 49 (*Bryopsis* sp. without LM) or 37 (*Bryopsis* sp. with LM) biological replicates.

**Fig. S3.**
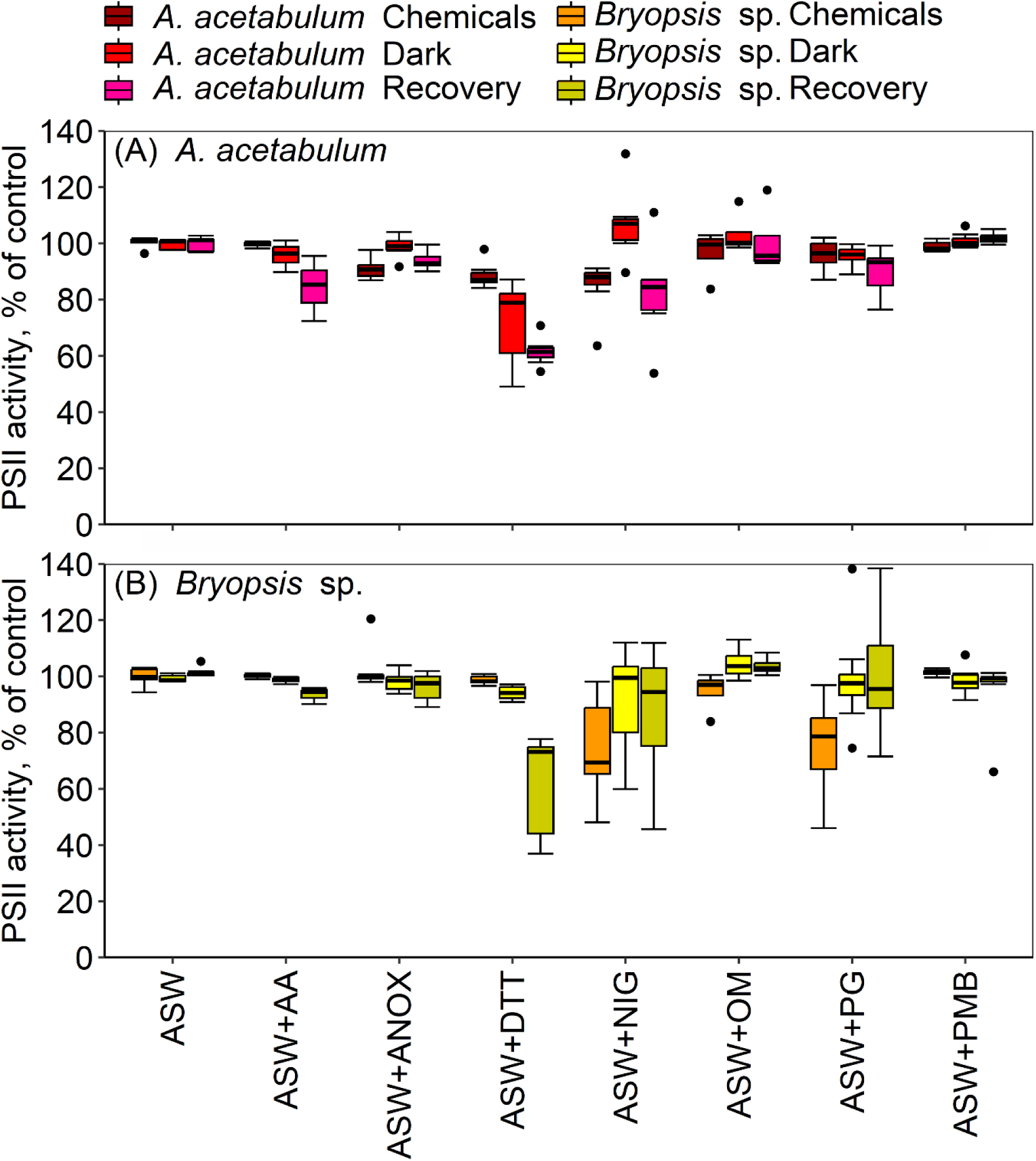
Effects of different chemicals on PSII activity in darkness in *Acetabularia acetabulum* (A) and *Bryopsis* sp. (B). The algae were incubated in the dark with the chemicals for 20 min (Chemicals) or 90 min (Dark), after which they were let to recover at low light (PPFD of 10–20 µmol m^-2^ s^-1^) for two h and again dark-acclimated for 20 min (Recovery). All the treatments were conducted at room temperature in artificial seawater (ASW) supplemented with the following chemicals: AA (antimycin A), anoxia (ANOX; glucose, glucose oxidase and catalase; to induce anaerobicity; Fig. S4), DTT (dithiothreitol), NIG (nigericin), OM (oligomycin), PG (propyl gallate) and PMB (polymyxin B). PSII activity was quantified with the chlorophyll *a* fluorescence parameter F_V_/F_M_. Control refers to the F_V_/F_M_ values of each sample prior to the addition of the chemicals (Chemicals) or after addition of the chemicals (Dark and Recovery). The box plots show medians, 2^nd^ and 3^rd^ quartiles, error bars show minimum and maximum values and dots show outliers (> 1.5 times the interquartile range), calculated based on three to eight biological replicates.

**Fig. S4.**
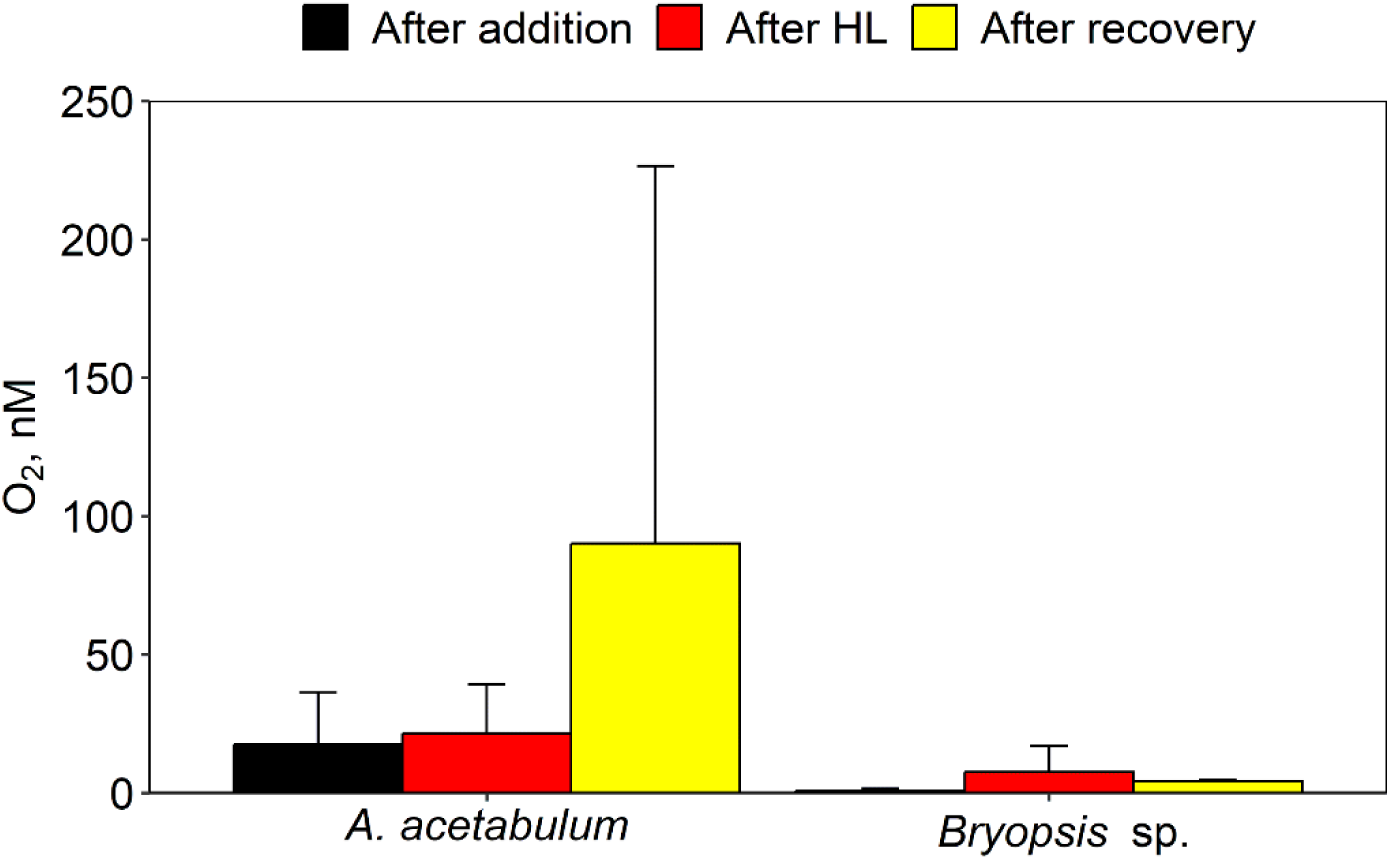
Oxygen concentration in *Acetabularia acetabulum* and *Bryopsis* sp. samples kept in two mL of artificial seawater in the presence of glucose, glucose oxidase and catalase, measured right after addition of the chemicals (black bars), after 50 min illumination with constant white high light (HL; red bars; PPFD 500 µmol m^-2^ s^-1^) and after subsequent two h at low light (yellow bars; PPFD 10–20 µmol m^-2^ s^-1^). All the treatments were conducted at room temperature. The bars show averages and error bars standard deviations calculated based on three to four biological replicates.

**Fig S5.**
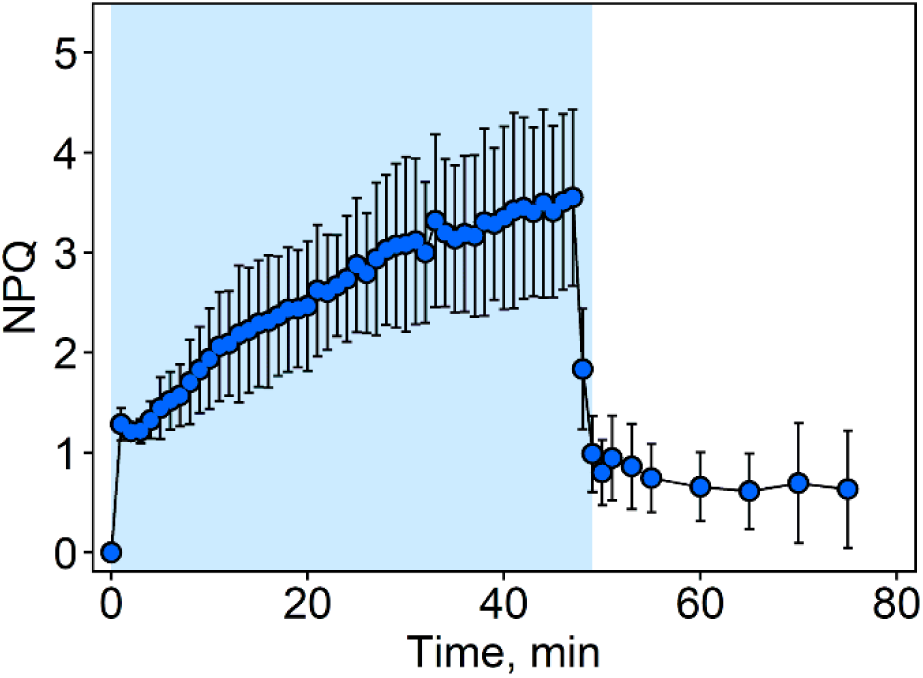
Non-photochemical quenching (NPQ), measured with chlorophyll *a* fluorescence as (F_M_-F_M_’)/F_M_’, in *Acetabularia acetabulum.* Dark acclimated algae were illuminated for 50 min with constant blue light (illustrated with the blue panel; PPFD 500 µmol m^-2^ s^-1^) and subsequently incubated in the dark for 25 min. During the last two minutes of the illumination, light intensity was gradually lowered to zero. The treatment was conducted at room temperature in artificial seawater. Symbols show averages and error bars standard deviations, calculated based on four biological replicates.

**Fig S6.**
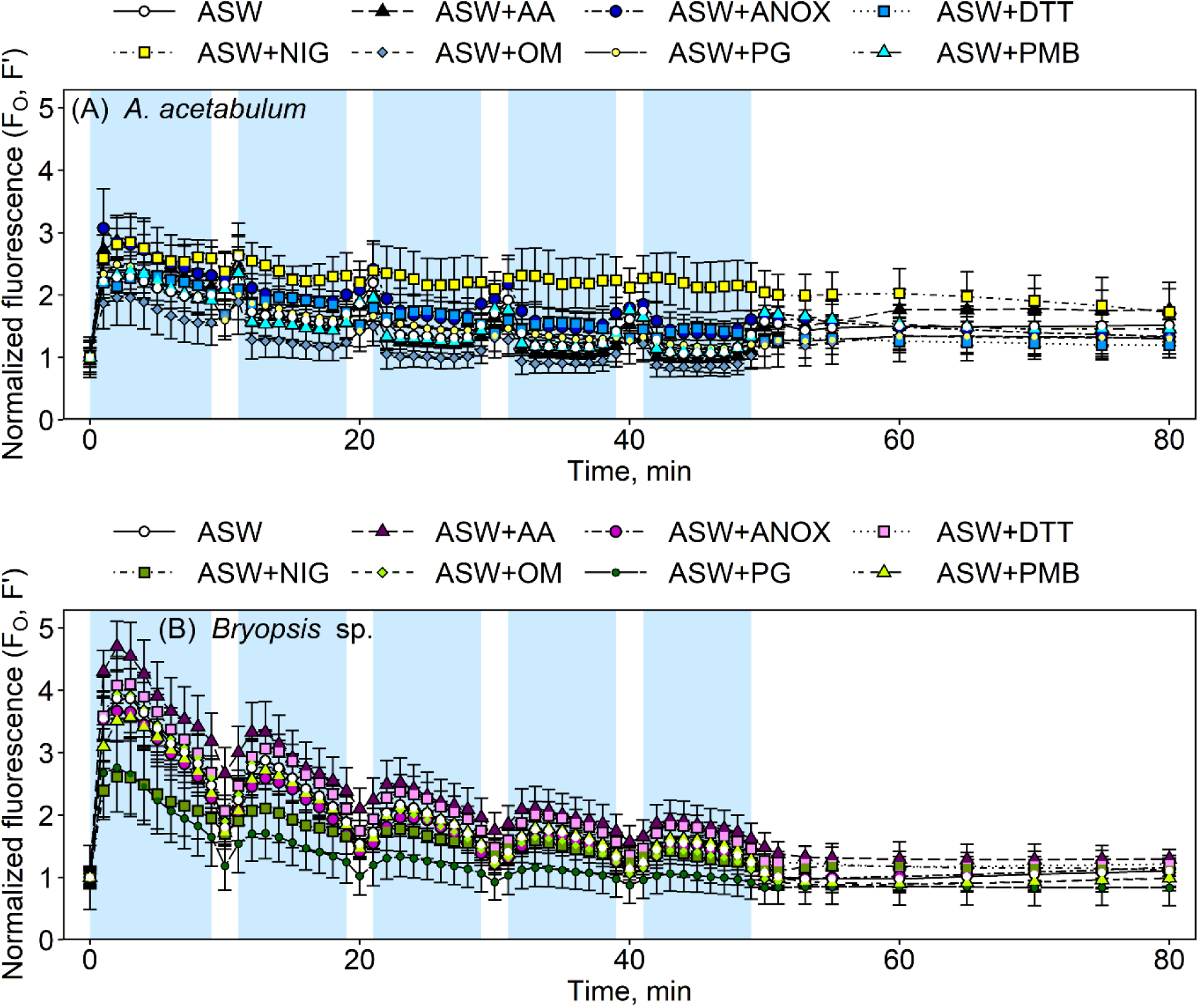
Chlorophyll *a* fluorescence yield (F_O_, F’; normalized to the starting values) in dark-acclimated *Acetabularia acetabulum* (A) and *Bryopsis* sp. (B) during a 50-min illumination (illustrated with the blue panels) with gradually fluctuating blue light (PPFD of 0 to 1000 µmol m^-2^ s^-1^; Fig. S1) and a subsequent 30-min darkness. The treatments were conducted at room temperature in artificial seawater (ASW) supplemented with the following chemicals: AA (Antimycin A), ANOX (glucose, glucose oxidase and catalase; to induce anaerobicity; Fig. S4), DTT (dithiothreitol), NIG (nigericin), OM (oligomycin), PG (propyl gallate) or PMB (polymyxin B), as indicated. Symbols show averages and error bars standard deviations, calculated based on four to 12 biological replicates.

**Fig. S7.**
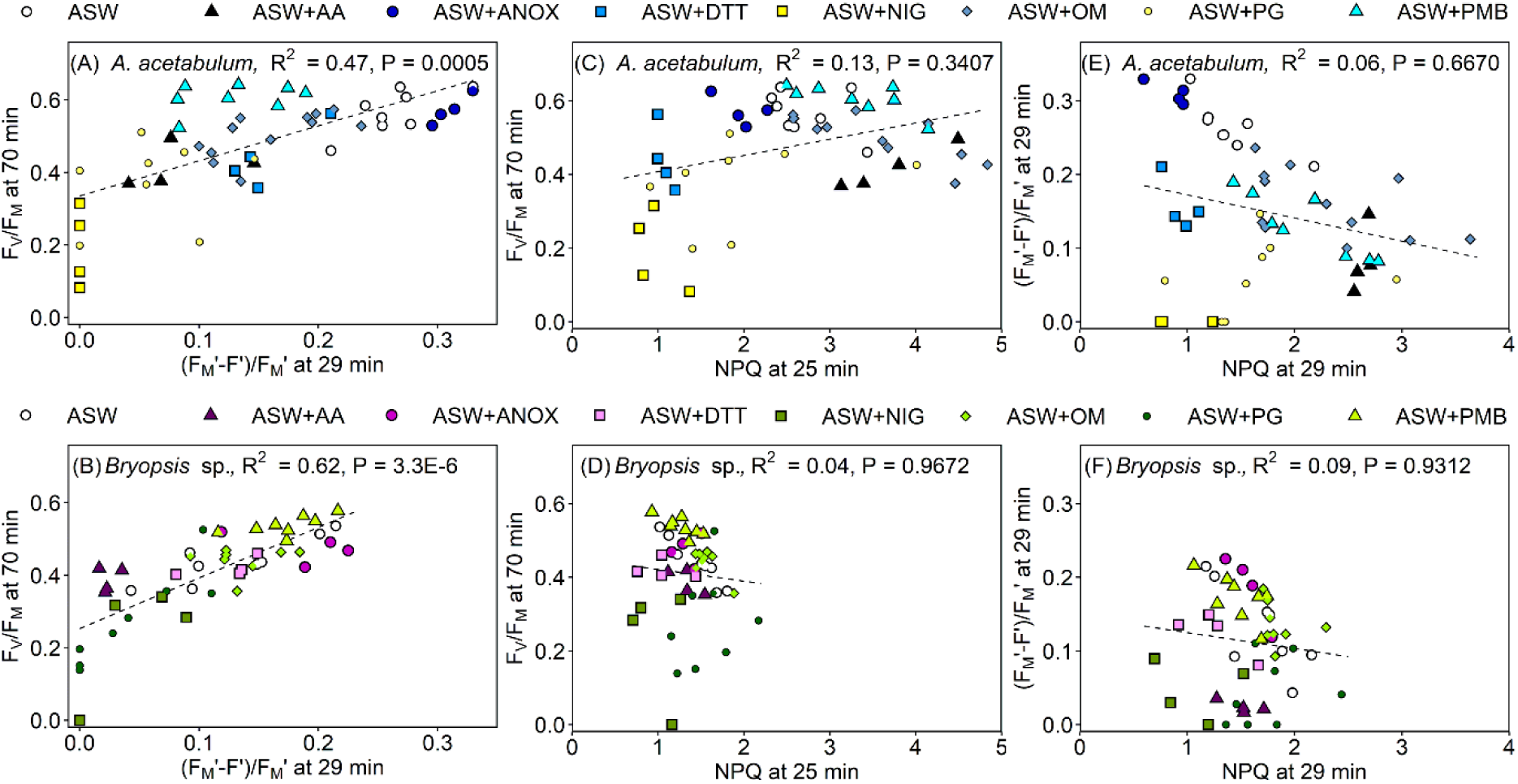
Correlations among PSII photoinhibition, PSII activity in light ((F_M_’-F’)/F_M_’) and non-photochemical quenching (NPQ) in *Acetabularia acetabulum* (A, C and E) and *Bryopsis* sp. (B, D and F). Photoinhibition was probed by the chlorophyll *a* fluorescence parameter F_V_/F_M_ after a 50-min fluctuating light treatment (blue light of PPFD 0 to 1000 µmol m^-2^ s^-1^; Fig. S1) and subsequent 20-min darkness (i.e., at 70 min). (F_M_-F’)/F_M_’ and NPQ, quantified as (F_M_-F_M_’)/F_M_’, were calculated at the indicated time points (at 29 min, the PPFD was ∼150 µmol m^-2^ s^-1^ and at 25 min, 1000 µmol m^-2^ s^-1^). For the original fluorescence traces, see Figs 5 and S6. All the treatments were conducted at room temperature in artificial seawater (ASW) supplemented with the following chemicals: AA (Antimycin A), ANOX (glucose, glucose oxidase and catalase; to induce anaerobicity; Fig. S4), DTT (dithiothreitol), NIG (nigericin), OM (oligomycin), PG (propyl gallate) or PMB (polymyxin B), as indicated. The dashed lines show best fits to a linear equation (statistics are highlighted in the corresponding panels). Symbols show individual biological replicates.

**Fig. S8.**
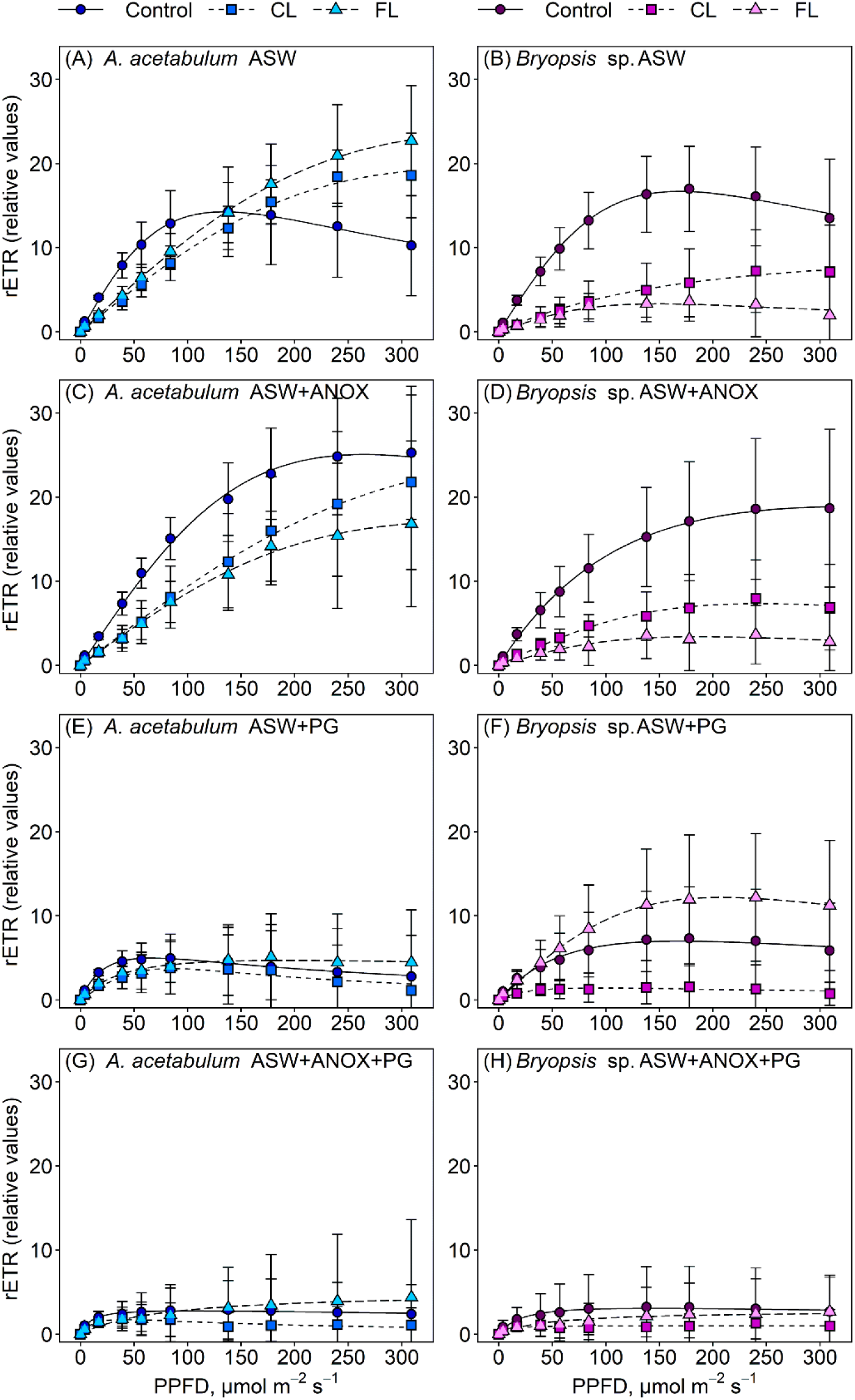
Rapid light response curves, measured with chlorophyll *a* fluorescence from light-acclimated *Acetabularia acetabulum* and *Bryopsis* sp., either before (circles; Control) or after a 50-min illumination at room temperature in artificial seawater (ASW) with constant (squares; CL; PPFD 500 µmol m^-2^ s^-1^) or fluctuating (triangles; FL; PPFD of 0 to 1000 µmol m^-2^ s^-1^) white light. During the light curve measurements, algae were kept in ASW (A, B), supplemented with glucose, glucose oxidase and catalase (C, D; ANOX; to induce anaerobicity; Fig. 4), propyl gallate (G, F; PG) or both (E, F). The algae were illuminated for 60 s with increasing intensities of blue light (as indicated), after which a saturating pulse was fired to calculate relative rates of electron transfer (rETR), as (F_M_’-F’)/F_M_’ x 0.5 x 0.84 x PPFD. Symbols show averages and error bars standard deviations calculated based on eight to 16 biological replications. Lines show best fits to the light response curves, modelled according to Eilers and Peeters (1988).

## Supplementary tables

**Table S1.** Statistical significances of differences in photoinhibition (without and with lincomycin) and recovery between *Acetabularia acetabulum* illuminated under constant light in plain artificial sea water and the indicated treatments. See Fig. 1 for the data. The significances have been obtained with heteroscedastic t-tests, using Bonferroni corrections.

| Comparisons | Photoinhibition | Recovery | Photoinhibition +LM |
| --- | --- | --- | --- |
| ASW vs chemicals (CL): |  |  |  |
| ASW vs ASW + AA | 0.0291 | **0.0004\*\*** | 0.7717 |
| ASW vs ASW + ANOX | **0.0014\*** | **2.7E-8\*\*\*** | 0.1096 |
| ASW vs ASW + DTT | 0.3599 | **6.6E-15\*\*\*** | 0.0976 |
| ASW vs ASW + NIG | **3.6E-9\*\*\*** | **0.0014\*** | **0.0016\*** |
| ASW vs ASW + OM | 0.1091 | 0.4665 | 0.0071* |
| ASW vs ASW + PG | 0.8374 | **0.0028\*** | 0.0764 |
| ASW vs ASW + PMB | 0.2078 | 0.6373 | 0.6310 |
| No LM vs LM (ASW, CL) | 0.7982 | n.m. | - |
| CL vs FL (ASW) | 0.0910 | 0.3096 | 0.0580 |
| *A. acetabulum* vs *Bryopsis* sp. (ASW, CL) | 0.2399 | **2.4E-8\*\*\*** | 0.1237 |

\* = P<0.1 (Bonferroni corrected value for 10 comparisons 0.01); \* = P<0.05 (Bonferroni corrected value for 10 comparisons 0.005); \*\* = P<0.01 (Bonferroni corrected value for 10 comparisons 0.001); \*\*\* = P<0.001 (Bonferroni corrected value for 10 comparisons 0.0001). ASW = artificial sea water, AA = antimycin A, ANOX = anaerobicity, DTT = dithiothreitol, NIG = nigericin, OM = oligomycin, PG = propyl gallate, PMB = polymyxin B, LM = lincomycin, CL = constant light, FL = fluctuating light, n.m. = not measured.

**Table S2.** Statistical significances of differences in photoinhibition (without and with lincomycin) and recovery between *Acetabularia acetabulum* illuminated under fluctuating light in plain artificial sea water and the indicated treatments. See Fig. 1 for the data. The significances have been obtained with heteroscedastic t-tests, using Bonferroni corrections.

| Comparisons | Photoinhibition | Recovery | Photoinhibition +LM |
| --- | --- | --- | --- |
| ASW vs chemicals (FL): |  |  |  |
| ASW vs ASW + AA | 0.0152 | **0.001\*\*** | 0.5796 |
| ASW vs ASW + ANOX | **0.0045\*** | **9.0E-6\*\*\*** | **0.0008\*\*** |
| ASW vs ASW + DTT | 0.0588 | **3.1E-13\*\*\*** | 0.0096* |
| ASW vs ASW + NIG | **0.0001\*\*** | 0.0867 | **3.2E-6\*\*\*** |
| ASW vs ASW + OM | 0.5626 | 0.2221 | 0.9543 |
| ASW vs ASW + PG | 0.5065 | **0.0013\*** | 0.1655 |
| ASW vs ASW + PMB | 0.5229 | 0.7528 | 0.6849 |
| No LM vs LM (ASW, FL) | **0.0022\*** | n.m. | - |
| CL vs FL (ASW) | See Table S1 | See Table S1 | See Table S1 |
| *A. acetabulum* vs *Bryopsis* sp. (ASW, FL) | 0.3297 | **2.2E-5\*\*\*** | 0.0580 |

\* = P<0.1 (Bonferroni corrected value for 10 comparisons 0.01); \* = P<0.05 (Bonferroni corrected value for 10 comparisons 0.005); \*\* = P<0.01 (Bonferroni corrected value for 10 comparisons 0.001); \*\*\* = P<0.001 (Bonferroni corrected value for 10 comparisons 0.0001). ASW = artificial sea water, AA = antimycin A, ANOX = anaerobicity, DTT = dithiothreitol, NIG = nigericin, OM = oligomycin, PG = propyl gallate, PMB = polymyxin B, LM = lincomycin, CL = constant light, FL = fluctuating light, n.m. = not measured.

**Table S3.** Statistical significances of differences in photoinhibition (without and with lincomycin) and recovery between *Bryopsis* sp. illuminated under constant light in plain artificial sea water and the indicated treatments. See Fig. 1 for the data. The significances have been obtained with heteroscedastic t-tests, using Bonferroni corrections.

| Comparisons | Photoinhibition | Recovery | Photoinhibition +LM |
| --- | --- | --- | --- |
| ASW vs chemicals (CL): |  |  |  |
| ASW vs ASW + AA | 0.6301 | 0.1274 | 0.3987 |
| ASW vs ASW + ANOX | 0.0199 | 0.7495 | **0.0038\*** |
| ASW vs ASW + DTT | **0.0023\*** | **4.4E-7\*\*\*** | 0.1080 |
| ASW vs ASW + NIG | 0.8649 | 0.3398 | 0.1530 |
| ASW vs ASW + OM | **0.0017\*** | 0.1362 | 0.9947 |
| ASW vs ASW + PG | 0.0400 | 0.2980 | 0.0076* |
| ASW vs ASW + PMB | 0.0054* | 0.5795 | 0.0389 |
| No LM vs LM (ASW, CL) | 0.0106 | n.m. | - |
| CL vs FL (ASW) | 0.6581 | 0.0328 | 0.0450 |
| *A. acetabulum* vs *Bryopsis* sp. (ASW, CL) | See Table S1 | **See Table S1\*\*\*** | See Table S1 |

\* = P<0.1 (Bonferroni corrected value for 10 comparisons 0.01); \* = P<0.05 (Bonferroni corrected value for 10 comparisons 0.005); \*\* = P<0.01 (Bonferroni corrected value for 10 comparisons 0.001); \*\*\* = P<0.001 (Bonferroni corrected value for 10 comparisons 0.0001). ASW = artificial sea water, AA = antimycin A, ANOX = anaerobicity, DTT = dithiothreitol, NIG = nigericin, OM = oligomycin, PG = propyl gallate, PMB = polymyxin B, LM = lincomycin, CL = constant light, FL = fluctuating light, n.m. = not measured.

**Table S4.** Statistical significances of differences in photoinhibition (without and with lincomycin) and recovery between *Bryopsis* sp. illuminated under fluctuating light in plain artificial sea water and the indicated treatments. See Fig. 1 for the data. The significances have been obtained with heteroscedastic t-tests, using Bonferroni corrections.

| Comparisons | Photoinhibition | Recovery | Photoinhibition +LM |
| --- | --- | --- | --- |
| ASW vs chemicals (FL): |  |  |  |
| ASW vs ASW + AA | 0.1668 | 0.9820 | 0.3842 |
| ASW vs ASW + ANOX | **0.0003\*\*** | 0.1700 | 0.0265 |
| ASW vs ASW + DTT | 0.2745 | **8.0E-11\*\*\*** | 0.9713 |
| ASW vs ASW + NIG | 0.6760 | 0.0092* | 0.1061 |
| ASW vs ASW + OM | 0.0129 | 0.2966 | 0.4023 |
| ASW vs ASW + PG | **0.0044\*** | 0.3726 | **3.6E-5\*\*\*** |
| ASW vs ASW + PMB | 0.8475 | 0.8475 | 0.0266 |
| No LM vs LM (ASW, FL) | 0.6732 | n.m. | - |
| CL vs FL (ASW) | See Table S3 | See Table S3 | See Table S3 |
| *A. acetabulum* vs *Bryopsis* sp. (ASW, FL) | See Table S2 | **See Table S2\*\*** | See Table S2 |

\* = P<0.1 (Bonferroni corrected value for 10 comparisons 0.01); \* = P<0.05 (Bonferroni corrected value for 10 comparisons 0.005); \*\* = P<0.01 (Bonferroni corrected value for 10 comparisons 0.001); \*\*\* = P<0.001 (Bonferroni corrected value for 10 comparisons 0.0001). ASW = artificial sea water, AA = antimycin A, ANOX = anaerobicity, DTT = dithiothreitol, NIG = nigericin, OM = oligomycin, PG = propyl gallate, PMB = polymyxin B, LM = lincomycin, CL = constant light, FL = fluctuating light, n.m. = not measured.

**Table S5.** Statistical significances of differences in (F_M_’-F’)/F_M_’ and NPQ changes (from 51 to 53 min at the dark) between *Acetabularia acetabulum* illuminated under fluctuating light in plain artificial sea water and the indicated treatments. See Fig. 5 for the data. The significances have been obtained with heteroscedastic t-tests, using Bonferroni corrections.

| Comparisons | Change in $(F_M' - F')/F_M'$ | Change in NPQ |
| --- | --- | --- |
| ASW vs ASW + AA | 0.6755 | 0.9502 |
| ASW vs ASW + ANOX | 0.4977 | 0.4809 |
| ASW vs ASW + DTT | 0.6870 | **1.3E-5\*\*\*** |
| ASW vs ASW + NIG | 0.1435 | **0.0008\*\*** |
| ASW vs ASW + OM | **0.0070\*** | 0.0188 |
| ASW vs ASW + PG | 0.9445 | 0.0354 |
| ASW vs ASW + PMB | **0.0069\*** | **6.6E-5\*\*\*** |

\* = $P < 0.1$ (Bonferroni corrected value for 7 comparisons 0.0143); \* = $P < 0.05$ (Bonferroni corrected value for 7 comparisons 0.0071); \*\* = $P < 0.01$ (Bonferroni corrected value for 7 comparisons 0.0014); \*\*\* = $P < 0.001$ (Bonferroni corrected value for 7 comparisons 0.00014). ASW = artificial sea water, AA = Antimycin A, ANOX = anaerobicity, DTT = dithiothreitol, NIG = nigericin, OM = oligomycin, PG = propyl gallate, PMB = polymyxin B.

**Table S6.** Statistical significances of differences in rapid light response curve parameters between non-illuminated and illuminated *Acetabularia acetabulum* and between the absence and presence of the indicated chemicals. See Figs 6 and S8 for the data. The significances have been obtained with heteroscedastic t-tests, using Bonferroni corrections.

| Comparisons | Alpha | rETR<sub>MAX</sub> | I<sub>K</sub> |
| --- | --- | --- | --- |
| ASW: |  |  |  |
| CTL vs CL | **9.0E-8\*\*\*** | 0.0436 | 0.0069* |
| CTL vs FL | **3.3E-7\*\*\*** | 0.0089* | **2.5E-6\*\*\*** |
| ASW + ANOX: |  |  |  |
| CTL vs CL | **1.0E-8\*\*\*** | 0.8511 | 0.0729 |
| CTL vs FL | **7.2E-9\*\*\*** | 0.2124 | 0.0658 |
| ASW + PG: |  |  |  |
| CTL vs CL | **8.3E-6\*\*\*** | 0.3881 | 0.3557 |
| CTL vs FL | **5.9E-6\*\*\*** | 0.5161 | 0.1082 |
| ASW + PG + ANOX: |  |  |  |
| CTL vs CL | 0.0133 | 0.3221 | 0.3248 |
| CTL vs FL | **0.0014\*** | 0.5107 | 0.2228 |
| ASW vs ASW + ANOX (CTL) | 0.5858 | **1.5E-5\*\*\*** | **7.6E-5** |
| ASW vs ASW + PG (CTL) | 0.0498 | **0.0001\*\*** | **1.1E-6** |
| ASW vs ASW + PG + ANOX (CTL) | 0.6456 | **5.1E-7\*\*\*** | **0.0023\*** |

\* = $P < 0.1$ (Bonferroni corrected value for 12 comparisons 0.0091); \* = $P < 0.05$ (Bonferroni corrected value for 12 comparisons 0.0045); \*\* = $P < 0.01$ (Bonferroni corrected value for 12 comparisons 0.0009); \*\*\* = $P < 0.001$ (Bonferroni corrected value for 12 comparisons 9.1E-5). ASW = artificial sea water, ANOX = anaerobicity, PG = propyl gallate, CL = constant light, FL = fluctuating light, alpha = initial slope of a light response curve, rETR<sub>MAX</sub> = a maximum rate of the electron transfer (relative values), I<sub>K</sub> = saturating light intensity.

**Table S7.** Statistical significances of differences in rapid light curve parameters between non-illuminated and illuminated *Bryopsis* sp. and between in the absence and presence of the indicated chemicals. See Figs 6 and S8 for the data. The significances have been obtained with heteroscedastic t-tests, using Bonferroni corrections.

| Comparisons | Alpha | rETR<sub>MAX</sub> | I<sub>K</sub> |
| --- | --- | --- | --- |
| ASW: |  |  |  |
| CTL vs CL | **4.0E-8\*\*\*** | 0.0915 | 0.1694 |
| CTL vs FL | **1.8E-9\*\*\*** | **2.1E-5\*\*\*** | 0.4865 |
| ASW + ANOX: |  |  |  |
| CTL vs CL | **3.4E-6\*\*\*** | **0.0002\*\*** | 0.9535 |
| CTL vs FL | **1.7E-6\*\*\*** | **9.6E-6\*\*\*** | 0.0211 |
| ASW + PG: |  |  |  |
| CTL vs CL | 0.0245 | **0.0008\*\*** | 0.365 |
| CTL vs FL | **0.0025\*** | 0.1627 | 0.0153 |
| ASW + PG + ANOX: |  |  |  |
| CTL vs CL | 0.0187 | 0.1134 | 0.5331 |
| CTL vs FL | 0.0149 | 0.5722 | 0.7490 |
| ASW vs ASW + ANOX (CTL) | 0.6727 | 0.1251 | 0.0994 |
| ASW vs ASW + PG (CTL) | 0.2963 | **4.9E-5\*\*\*** | **0.0005\*\*** |
| ASW vs ASW + PG + ANOX (CTL) | 0.6665 | **8.5E-8\*\*\*** | 0.0179 |

\* = P<0.1 (Bonferroni corrected value for 11 comparisons 0.0091); \* = P<0.05 (Bonferroni corrected value for 11 comparisons 0.0045); \*\* = P<0.01 (Bonferroni corrected value for 11 comparisons 0.0009); \*\*\* = P<0.001 (Bonferroni corrected value for 11 comparisons 9.1E-5). ASW = artificial sea water, ANOX = anaerobicity, PG = propyl gallate, CL = constant light, FL = fluctuating light, alpha = initial slope of a light response curve, ETR<sub>MAX</sub> = a maximum rate of the electron transfer (relative values), I<sub>K</sub> = saturating light intensity.

